# Preclinical evaluation of the novel [^18^F]CHDI-650 PET ligand for non-invasive quantification of mutant huntingtin aggregates in Huntington’s disease

**DOI:** 10.1101/2024.05.16.594492

**Authors:** Franziska Zajicek, Jeroen Verhaeghe, Stef De Lombaerde, Annemie Van Eetveldt, Alan Miranda, Ignacio Munoz-Sanjuan, Celia Dominguez, Vinod Khetarpal, Jonathan Bard, Longbin Liu, Steven Staelens, Daniele Bertoglio

## Abstract

**Purpose:** Positron emission tomography (PET) imaging of mutant huntingtin (mHTT) aggregates is a potential tool to monitor disease progression as well as the efficacy of candidate therapeutic interventions for Huntington’s disease (HD). To date, the focus has been mainly on the investigation of ^11^C radioligands; however, favourable ^18^F radiotracers will facilitate future clinical translation. This work aimed at characterising the novel [^18^F]CHDI-650 PET radiotracer using a combination of *in vivo* and *in vitro* approaches in a mouse model of HD.

**Methods:** After characterising [^18^F]CHDI-650 using *in vitro* autoradiography, we assessed *in vivo* plasma and brain radiotracer stability as well as kinetics through dynamic PET imaging in the heterozygous (HET) zQ175DN mouse model of HD and wild-type (WT) littermates at 9 months of age. Additionally, we performed a head-to-head comparison study at 3 months with the previously published [^11^C]CHDI-180R radioligand.

**Results:** Plasma and brain radiometabolite profiles indicated a suitable metabolic profile for *in vivo* imaging of [^18^F]CHDI-650. Both *in vitro* autoradiography and *in vivo* [^18^F]CHDI-650 PET imaging at 9 months of age demonstrated a significant genotype effect (*p*<0.0001) despite the poor test-retest reliability. [^18^F]CHDI-650 PET imaging at 3 months of age displayed higher differentiation between genotypes when compared to [^11^C]CHDI-180R.

**Conclusion:** Overall, [^18^F]CHDI-650 allows for discrimination between HET and WT zQ175DN mice at 9 and 3 months of age. [^18^F]CHDI-650 represents the first suitable ^18^F radioligand to image mHTT aggregates in mice and its clinical evaluation is underway.

## Introduction

Huntington’s disease (HD) is a progressive autosomal dominant neurodegenerative disorder linked to the huntingtin (HTT) gene encoding for mutant huntingtin protein (mHTT), the latter containing an expanded polyglutamine (polyQ) repeat that leads to the formation of mHTT aggregates [1, 2]. Accumulation of mHTT is involved in cellular dysfunction and loss of neurons resulting in progressive motor, psychiatric, and cognitive impairments in people with HD (PwHD) [3].

Several therapeutic interventional strategies directed at mHTT-lowering are currently under clinical evaluation [4, 5]. Guiding the therapeutic development and monitoring disease progression in (pre-)clinical settings are therefore of great value. Dynamic positron emission tomography (PET) imaging of mHTT aggregates represents a strategic evaluation tool due to its brain region-based quantification compared to mHTT detection in non-brain compartments, such as CSF or plasma quantification using ELISA-based assays [6–8].

To date, we have described two ^11^C radiotracers for use in mHTT aggregate imaging, namely [^11^C]CHDI-626 and [^11^C]CHDI-180R [9–14]. Specifically, [^11^C]CHDI-626 displayed rapid clearance and kinetics that translated to limited utility for mHTT PET imaging in mice; nevertheless, it displayed significantly increased striatal binding in 3-month-old zQ175DN heterozygous (HET) mice compared to their wildtype (WT) littermates [12]. In contrast, [^11^C]CHDI-180R showed high binding potency to mHTT aggregates and was metabolically stable in mice [9, 11]. Longitudinal studies in HD mouse models suggest a high preclinical utility of [^11^C]CHDI-180R as a mHTT aggregate-directed PET radioligand in mouse [13] and nonhuman primate [14] models of HD. Nonetheless, the use of ^11^C radioligand is limited by the short half-life of the radioisotope (20.4 min). To further facilitate clinical translation and better clinical practicability, the longer half-life of ^18^F radionuclides (109.8 min) will be preferred, which will allow tracer produced at a cyclotron site to be transported to dispersed imaging facilities and allow better adjustment to scan duration and increased numbers of scans [15]. In addition, versatility of existing labelling process renders ^18^F radiotracers a cheaper, easier, and thus a more applicable option for PET imaging [16]. This consideration led to the synthesis of an ^18^F-based radioligand with good pharmacological characteristics, namely [^18^F]CHDI-961 [17]. [^18^F]CHDI-961, however, exhibited bone uptake *in vivo* likely due to fluorine-containing metabolites formed through oxidative defluorination, which negatively impacts brain imaging analysis. To suppress this oxidative defluorination, and consequently avoid accumulation of [^18^F]-signal in bones, deuterium atoms were incorporated at the proposed site for oxidation, resulting in an improved radioligand, [^18^F]CHDI-650 [17]. Using the zQ175DN mouse model of HD, we first compared the *in vivo* plasma and brain stability of [^18^F]CHDI-961 and [^18^F]CHDI-650. Further, we examined the *in vivo* kinetic properties of [^18^F]CHDI-650 to quantify mHTT aggregates, including test-retest reliability and detectability of genotype effect. Finally, we performed a head-to-head comparison with [^11^C]CHDI-180R [13] in 3-month-old zQ175DN mice to test [^18^F]CHDI-650 performance at an age with minimal mHTT aggregate load.

## Material and Methods

### Animals

A total of 131 9-month-old male HET (*n*=65) and WT (*n*=66), as well as 48 3-month-old male HET (*n*=24) and WT (*n*=24) zQ175DN knock-in mice (B6.129S1-Htt^tm1.1Mfc^/190ChdiJ JAX stock #029928) [18, 19], were obtained from Jackson Laboratories (Bar Harbor, Maine, USA). Due to sporadic congenital portosystemic shunt occurring in C57BL/6J mice [20], all animals were screened at Jackson Laboratories before shipment in order to avoid this confounding factor; hence all animals used in this study were shunt-free. Group size, body weight, and age of the animals are reported in Supplementary Table S1-S5. Animals were group-housed in ventilated cages under a 12 h light/dark cycle in a temperature- and humidity-controlled environment with food and water ad libitum. The animals were given at least one week for acclimatisation after arrival before the start of any procedure. For the ARG study, brains from 9-month-old male HET (*n*=10) zQ175DN knock-in mice, as well as age-matched WT littermates (*n*=10), were obtained from Jackson Laboratories (Bar Harbour, Maine, USA) (ID = CHDI-81003019). Experiments followed the European Committee (decree 2010/63/CEE) and were approved by the Ethical Committee for Animal Testing (ECD 2019-39) at the University of Antwerp (Belgium).

### Radioligand synthesis

[^18^F]CHDI-650 and [^18^F]CHDI-961 were synthesised as previously described [17] but adapted to an automated synthesis module (Trasis AllInOne, Belgium). The molecular structures for [^18^F]CHDI-961 and [^18^F]CHDI-650 are reported in Fig. S1. Firstly, [^18^F]fluoride was generated by cyclotron bombardment of an [^18^O]H_2_O target (typical starting activity: ±20 GBq). The [^18^F]fluoride was transported to the Trasis module and trapped on a Sep-Pak QMA cartridge (Waters, preconditioned with 5 ml 0,1 M K_2_CO_3_ and 5 ml ultrapure water). It was then eluted with 0.8 ml of 38 mM tetra-ethyl ammonium bicarbonate (TEAB) solution (9/1 AcN/water) in a reactor, followed by azeotropic drying. Afterwards, the Tosylate precursor (3 mg in 1 ml DMSO for [^18^F]CHDI-650; 5 mg in 1 ml DMF for [^18^F]CHDI-961) was added. The reaction was allowed to proceed at 100°C for 5 minutes.

For [^18^F]CHDI-650, the reaction mixture was quenched with 4 ml of ultrapure water and then transferred to the HPLC system using a Waters XBridge C18, 5µm, 10×150 mm column (Waters Belgium), eluted with ethanol / 0.05M sodium acetate (NaOAc) (pH5.5) (30/70; v/v) as mobile phase, at a flow rate of 3 ml/min that allowed for isolation of the desired [^18^F]CHDI-650 (typical collection volume: 5 ml). The collected peak was then diluted with 25 ml of ultrapure water. This solution was loaded on a Sep-Pak Light tC18 cartridge (Waters, preconditioned with 5 ml ethanol and 5 ml ultrapure water) and rinsed with 5 ml of ultrapure water. [^18^F]CHDI-650 was eluted with 1 ml of 50% ethanol in water, followed by dilution with 4 ml of saline. Finally, this solution was sterile-filtered with an in-line filter (Cathivex GV 25 mm disk size; 0.22 µm).

For [^18^F]CHDI-961, the reaction mixture, diluted with ultrapure water + 0.1% ascorbic acid was purified on an HPLC system using a Kinetex EVO C18, 100 Å, 10×250 mm column (Phenomenex), eluted with ethanol / 0.05M NaOAc (pH5.5 + 0.01% ascorbic acid) (30/70; v/v) as mobile phase, at a flow rate of 3 ml/min that allowed for isolation of the desired [^18^F]CHDI-961 (typical collection volume: 5 ml). The collected peak was then diluted with 10ml of ultrapure water + 0.1% ascorbic acid. This solution was passed through a Sep-Pak Light tC18 cartridge (Waters, preconditioned with 5 ml ethanol and 5 ml ultrapure water) and washed with 5 ml of ultrapure water. [^18^F]CHDI-961 was released with 0.5 ml of 96% ethanol via a Nalgene sterilising filter (Thermo Scientific; 0.2 µm) and diluted with 5 ml of saline.

Afterwards, for both [^18^F]CHDI-650 and [^18^F]CHDI-961 a Luna C18 (2), 5μm, 100 Å∼ 4.6 mm x 250 mm (Phenomenex) HPLC column was used, with acetonitrile/0.05 M NaOAc pH 5.5 (60:40 for 10 min v/v, followed by a gradient to 10:90 and reverse for 10 min) as mobile phase, at a flow of 1 ml/min, with UV absorbance set at 283 nm. For [^18^F]CHDI-650, the radiochemical purity determined was greater than 99% and molar activity was 223.55±67.57 GBq/µmol (*n*=30). The decay-corrected radiochemical yield was 25.14±10.20%. For [^18^F]CHDI-961, the radiochemical purity was greater than 99% and molar activity was 278.54±56.19 GBq/µmol (*n*=6). The decay-corrected radiochemical yield was 5.45±2.23%.

For the comparison study in 3-month-old zQ175DN mice, [^11^C]CHDI-180R was synthesised as previously described [13]. Radiochemical purity was greater than 99% and the molar activity was 58.91±9.89 GBq/µmol (*n*=11). The decay-corrected radiochemical yield was 13.36±1.91%.

### Radiometabolite analysis

Evaluation of *in vivo* plasma and brain radiometabolite profiles was performed at 5, 15, 30, 45, 60, and 90 min post-injection (p.i.) for [^18^F]CHDI-650 and at 5, 15, 25, and 45 min p.i. for [^18^F]CHDI-961. The number of animals is reported in the supplementary Table S4. Mice were injected via the lateral tail vein with [^18^F]CHDI-650 (WT: 6.4±1.4 MBq; HET: 6.4±1.3 MBq in 200 μl) or [^18^F]CHDI-961 (WT: 8.6 ± 0.8 MBq; HET: 9.1 ± 0.8 MBq in 200 μl) and blood was withdrawn via cardiac puncture at designated times and brains were immediately dissected. Plasma samples (150 μl), obtained after centrifugation of blood at 4500×rpm (2377xrcf) for 5 min, were mixed with equal amounts of ice-cold acetonitrile. Whole blood and plasma were counted on the gamma counter to obtain the plasma-to-whole blood ratio (p/b ratio). Then, 10 µl of non-labelled reference was added to plasma samples (1 mg/ml) and subsequently centrifuged at 4500×rpm (2377xrcf) for 5 min to precipitate denatured proteins. The supernatant was separated from the precipitate and both fractions were counted in a gamma counter to calculate the extraction efficiency (percentage of recovery of radioactivity), which was 96.2±16.0% for [^18^F]CHDI-650 and 97.3±3.6% for [^18^F]CHDI-961. Brain samples were homogenised with 1 ml of ice-cold acetonitrile, 10 µl of non-labelled reference were added (1 mg/ml) and subsequently centrifuged at 4500×rpm (2377xrcf) for 5 min to precipitate denatured proteins. The supernatant was separated from the precipitate and both fractions were counted in a gamma counter to calculate the extraction efficiency, which was 97.9±2.8% for [^18^F]CHDI-650 and 95.2±7.0% for [^18^F]CHDI-961. Next, 100 μl of plasma or brain supernatant was loaded onto a pre-conditioned reverse-phase (RP)-HPLC system (Luna C18(2), 5 μm HPLC column (250×4.6 mm) + Phenomenex security guard pre-column) and eluted with NaOAc 0.05M pH 5.5 and acetonitrile: (60:40 for 10 min v/v, followed by a gradient to 10:90 and reverse) buffer at a flow rate of 1 ml/min. RP-HPLC fractions were collected at 0.5 min intervals for 10 min and radioactivity was measured in a gamma counter. The radioactivity present in each peak was expressed as a percentage of the total radioactivity eluted (i.e., the sum of radioactivity in all fractions) in counts per minute (CPM). Individual radioactivity in CPM were standardised by bodyweight and injected activity at the start of the gamma counter run. To determine the recovery of [^18^F]CHDI-650 as well as the stability of the tracer during the workup, experiments were performed using control blood spiked with 185 kBq of the radiotracer or by pipetting the latter directly into the brain homogenate. Sample workup, which was identical to the procedure described above, confirmed that no degradation of the tracers occurred during preparation in both plasma (99.2±0.0% intact [^18^F]CHDI-650; 99.6±0.0% intact [^18^F]CHDI-961) and brain (99.0±0.0% intact [^18^F]CHDI-650; 99.6±0.0% intact [^18^F]CHDI-961).

### Dynamic PET imaging

Dynamic microPET/Computed tomography (CT) images were acquired on Siemens Inveon PET-CT scanners (Siemens Preclinical Solution, Knoxville, USA). Animal preparation was performed as previously described [21, 22]. Briefly, the animals were placed side by side on the scanner bed with the heart of the animals in the scanner’s field of view. Anaesthesia was induced by inhalation of isoflurane (5% for induction, and 1.5-2% for maintenance during preparation and scanning) supplemented with oxygen. After induction, all mice were catheterised in the tail vein for intravenous (i.v.) bolus injection of the tracer and placed on the scanner bed. The respiration and heart rate of the animal were constantly monitored using the Monitoring Acquisition Module (Minerve, France) during the entire scanning period. The core body temperature of the animals was maintained using a warm airflow. At the onset of the dynamic microPET scan, mice were injected with a bolus of radiotracer over a 12-second interval (1 ml/min) using an automated pump (Pump 11 Elite, Harvard Apparatus, USA). Tracer was injected with activity as high as possible to obtain good image quality while keeping the cold dose as low as possible to minimise any potential mass effect. Animal and dosing information for both WT and HET mice of all paradigms are given in Table S1-S3.

PET data were acquired in list mode format. Following the microPET scan, a 10 min 80 kV/500 μA CT scan was performed for attenuation and scatter correction. During the same microPET scan, the blood radioactivity was estimated using a non-invasive image-derived input function (IDIF). The IDIF was extracted from the PET images by generating a heart volume-of-interest (VOI) on the early time frames, considering a lower threshold of 50% of the maximum activity as previously described [21–23]. Since the peripheral metabolic profile of [^18^F]CHDI-650 displayed an increasing radioactive uptake in the liver becoming noticeable around 60 min of scan, we applied an additional step in order to rule out potential spill-in from the liver in the VOI of the IDIF. Therefore, we averaged the activity from 60 to 90 min to further restrict the VOI of the IDIF by excluding those voxels with an average activity exceeding twice the minimal average activity within this time frame. The resulting VOI was used to extract the entire IDIF curve.

Acquired PET data were reconstructed into 33 (60 min) or 39 (90 min) frames of increasing length (12×10s, 3×20s, 3×30s, 3×60s, 3×150s, and 9×300s or 15×300s). Images were reconstructed using a proprietary list-mode iterative reconstruction with spatially variant resolution modelling in 8 iterations and 16 subsets of the 3D ordered subset expectation maximisation (OSEM 3D) algorithm [24]. Normalisation, dead time, and CT-based attenuation corrections were applied. PET image frames were reconstructed on a 128x128x159 grid with 0.776×0.776×0.796 mm^3^ voxels.

### Image processing and analysis

Image processing and analysis were performed as previously described with PMOD 3.6 software (PMOD Technologies, Zurich, Switzerland) [12]. For spatial normalisation of the PET/CT images, brain normalisation of the CT image to the CT template was done by adjusting the previously described procedure [25]. The VOIs of the Waxholm atlas [26] were also adjusted as in previous studies [12, 13] and time activity curves (TACs) of the different brain regions were then extracted. The IDIF was extracted from the PET images measuring the whole blood activity in the lumen of the left ventricle as described above. For [^18^F]CHDI-650 the whole-blood IDIF was corrected for radiometabolites and the p/b ratio was applied for subsequent kinetic modelling. For [^11^C]CHDI-180R, no metabolite correction was needed given the high stability *in vivo* (>95% at 40 min p.i.) nor p/b ratio correction given the stable profile. Thus the uncorrected IDIFs were used for further analyses as previously validated [13]. TACs, reported as standardised uptake value (SUV), were analysed with kinetic modelling to determine the total volume of distribution (*V*_T_) using IDIFs to estimate *V*_T(IDIF)_ as a surrogate of *V*_T_. Kinetic modelling was performed by fitting these TACs with Logan linear model to calculate *V*_T(IDIF)_. The linear phase was determined from the curve fitting based on 10% maximal error. Model fits for Logan were used based on qualitative assessment (i.e. adequate fit) and standard error percentage (SE%) (i.e. only if <20%). Animals and regions not passing both metrics were excluded from the analysis; animal numbers for each paradigm are listed in Table S5. Following models were investigated but no adequate fit could be established due to failure during qualitative assessment or SE%: 2-tissue compartment model (2TCM), 2TCM with coupled *K*_1_/*k*_2_ fit & *V*_T_ (2TCM *K*_1_/*k*_2_), 2TCM with vascular trapping (2TCM VascTrap), 3-tissue compartment model (3TCM), 3-sequential-TCM (3sTCM), and 3sTCM with *k*_6_ at 0 (3sTCM *k*_6_=0). These compartmental models vary in their estimated parameters used for both model curve fitting and *V*_T(IDIF)_ calculation, thus they represent a variety of modelling options. Parameters and their standard errors for each compartmental model are given in Table S6. Voxel-wise kinetic modelling was performed using the Logan graphical analysis in PXMOD (PMOD). The averaged parametric maps were subsequently overlaid onto a Waxholm MRI template for anatomical visualisation.

### Autoradiography

Procedure was performed as previously described [17]. Sections were air-dried at room temperature (RT) and pre-incubated for 20 min with binding buffer (50mM Tris-HCl + 120 mM NaCl + 5mM KCl + 2mM CaCl_2_ + 1mM MgCl_2_, pH 7,4). Incubation with total binding solution (0.005 to 3nM of [^18^F]CHDI-650 in binding buffer) and non-specific binding solution (0.005 to 3nM of [^18^F]CHDI-650 + 10 µM of cold CHDI-650 ligand in binding buffer) was performed for 1 h at RT. Subsequently, sections were washed three times for 10 min in ice-cold washing buffer (50mM Tris-HCl, pH 7,4), followed by a brief wash in ice-cold distilled water, and dried for 30 min at RT. For calibration, 2 µl of solutions ranging from 0.000625 nM to 5.12 nM were pipetted onto absorbent paper. Finally, all sections together with the calibration solution spots were exposed on imaging plates (BAS-IP2040, Fujifilm, Japan) for 10 min. Radioactivity was detected with a phosphor imager (Fuji FLA-7000 image reader) and quantified based on intensity values calculated using the calibration spots and fitted standard curves. The anatomical reference images shown in the results are the same as those previously used [13]. Analysis was done in Fiji, ImageJ (version: 2.1).

### Statistical analysis

The Shapiro-Wilk test revealed that *in vivo* data and *in vitro* data were normally distributed. Nonlinear fit was used for tracer signal calculation from autoradiography gray values. Rational fit and linear fit were used for IDIF correction. An unpaired T-test was performed to investigate differences between genotypes in dosing parameters for scan acquisition and during autoradiography analysis. A paired T-test was performed to investigate differences between test and retest in dosing parameters for scan acquisition. A Spearman’s correlation with interpolation (sigmoidal, 4PL, X is log_(concentration)_) was applied to injected masses and *V*_T(IDIF)_ values from Logan for visualisation of a potential mass dose effect. Regular two-way ANOVA with post hoc Bonferroni correction for multiple comparisons was applied to analyse *V*_T(IDIF)_ value differences between genotypes.

The individual relative test-retest variability was calculated as follows:

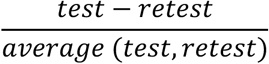

The individual absolute test-retest variability was calculated as follows:

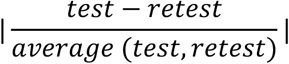

In addition, a Bland-Altman analysis was performed to assess the agreement between two measures on an individual level. All statistical analyses above were performed with GraphPad Prism (v9.2.0) statistical software. Furthermore, the Intraclass correlation coefficient (ICC) for the test-retest variability study, reported in the literature to classify the variability of tracers, was calculated with the software JMP Pro 17 (v17.0.0). Group x timepoint interaction was set as a fixed effect. Animal and timepoint as well as the interaction of animal x group and animal x timepoint were estimated as random effects. The interaction animal x group and animal x timepoint did not impact the ICC calculation. The ICC is reported as the ratio of the animal variance component for the effect to the sum of positive variance components. In case the animal variance component was estimated as negative the ICC is reported as 0. One-tailed power analyses were performed in G*Power (v3.1.9.6) with significance level (*α*) at 0.05 and confidence (*β*) set to 80%. Data are represented as mean ± standard deviation (SD) unless specified otherwise. All tests were two-tailed unless stated otherwise. Statistical significance was set at *p*<0.05 with the following standard abbreviations used to reference *p* values: ns, not significant; \**p*<0.05; \*\**p*<0.01; \*\*\**p*<0.001; \*\*\*\**p*<0.0001.

## Results

### [^18^F]CHDI-650 autoradiography in zQ175DN mice displays phenotypic difference

To confirm that [^18^F]CHDI-650 can differentiate between genotypes that differ in expression of mHTT aggregates within the brain, as previously demonstrated [17], an autoradiography study was performed (Fig. 1a). A significant difference between the genotypes was observed in both striatum and motor cortex (*p*<0.0001) (Fig. 1b; 1c). As we confirmed [^18^F]CHDI-650 differentiates between genotypes further *in vivo* analysis was conducted.

**Fig. 1.**
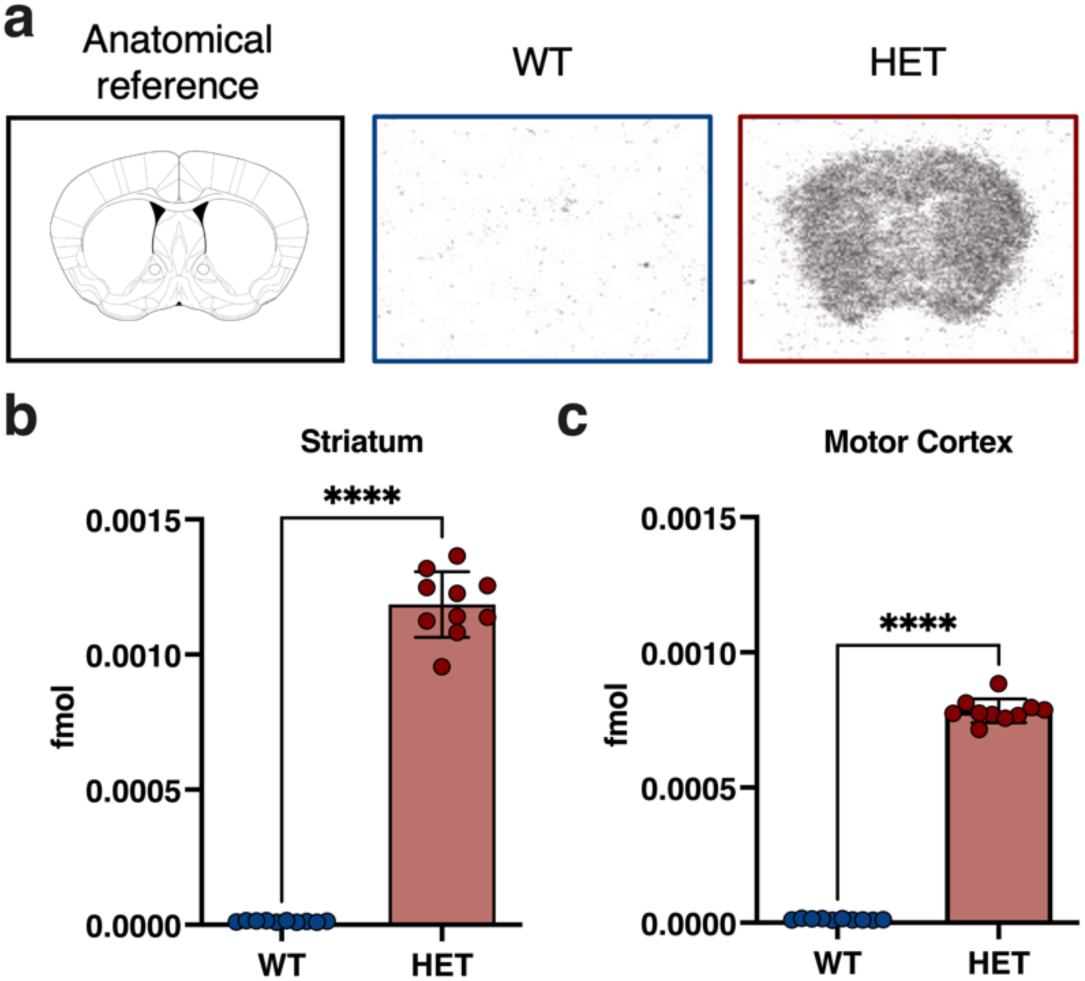
[^18^F]CHDI-650 autoradiography displays specific binding only in HET mice. a, Exemplary autoradiography images for a WT and a HET 9-month-old mouse. Quantification of mHTT aggregates in Striatum (b) and Motor Cortex (c). \*\*\*\**p*<0.0001. b and c: WT: *n*=10; HET: *n*=10

### [^18^F]CHDI-650 demonstrates improved plasma and brain stability compared to [^18^F]CHDI-961

[^18^F]CHDI-650 and [^18^F]CHDI-961 radiometabolite analysis was performed to determine the presence of radiometabolites over time. Comparison of the plasma radiometabolite profile of the two radiotracers revealed more intact radioligand and a slower elimination profile for [^18^F]CHDI-650 relative to [^18^F]CHDI-961. At 40 min p.i., 19.24±3.79% of total radioactivity in plasma was accounted for by intact [^18^F]CHDI-961 while for [^18^F]CHDI-650 at 45 min p.i., 30.41±5.08% of total radioactivity was due to the intact radioligand (Fig. 2a). The radiometabolite analysis of [^18^F]CHDI-650 (Fig. S2) and [^18^F]CHDI-961 (Fig. S3) demonstrated at least two highly polar radiometabolites in plasma of both genotypes. [^18^F]CHDI-650 displayed a similar plasma profile in WT and HET zQ175DN mice (Fig. S2); therefore, the combined radiometabolite data was used for generating a population-based metabolite-correction curve based on a rational fit (Fig. 2b) for correction of the extracted IDIFs. Next, a p/b ratio correction based on a linear fit (Fig. 2c) was also included to generate the final plasma metabolite-corrected IDIFs for kinetic modelling (Fig. 2d). Similar to the plasma profile, also the brain radiometabolite profile demonstrated the improved stability of [^18^F]CHDI-650 compared to [^18^F]CHDI-961. However, both radiotracers displayed differences in their brain profile in WT compared to HET zQ175DN mice (Fig. S2 and Fig. S3). At 40 min p.i. 26.92±6.07% and 76.50±4.27% of total radioactivity in brain was accounted for by intact [^18^F]CHDI-961 in WT and HET, respectively (Fig. S3). On the other hand, for [^18^F]CHDI-650 at 45 min p.i., 51.03±3.73% and 93.67±0.67% of total radioactivity was due to the intact radioligand for WT and HET, respectively (Fig. S2). Nonetheless, this difference was solely related to the specific uptake, thus longer retention time, in the brain of HET mice compared to WT mice as shown in the standardised CPM of [^18^F]CHDI-650 (Fig. S4). Accordingly, both genotypes displayed negligible radiometabolites in the brain over time, suggesting no brain-penetrating species.

**Fig. 2.**
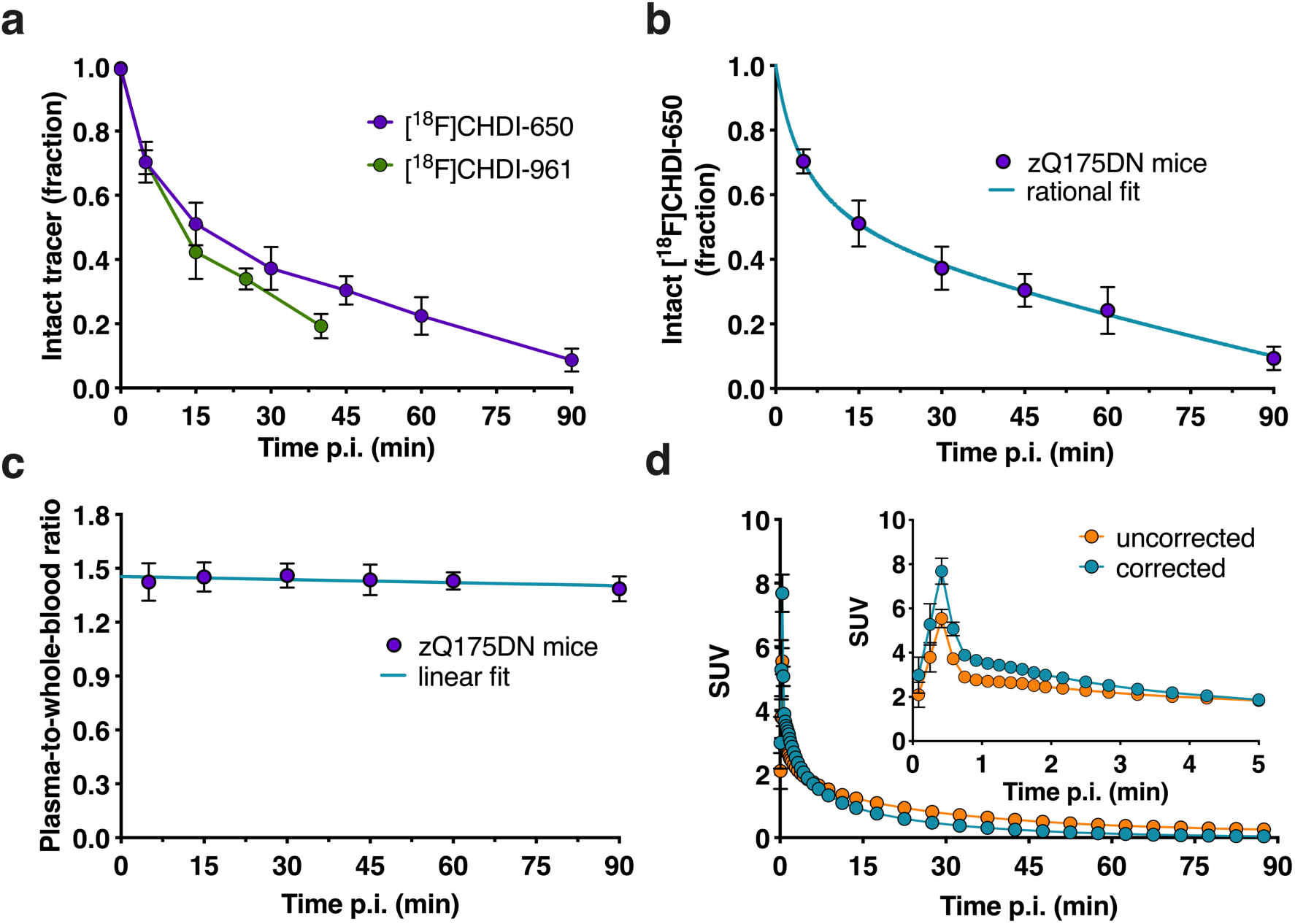
Blood analysis of [^18^F]CHDI-961 and [^18^F]CHDI-650. a, Comparison of [^18^F]CHDI-961 and [^18^F]CHDI-650 intact radioligand as fraction in plasma. b, Fraction of intact [^18^F]CHDI-650 in plasma with rational fit applied. c, Plasma-to-whole-blood ratio with linear fit. d, Averaged uncorrected and corrected IDIF SUV curves of [^18^F]CHDI-650 over 90 min scan time with magnification on x-axis. Data is based on the entire data (WT and HET zQ175DN mice). In d, data are represented as mean±SEM. a-c: *n*=2-6 per genotype, time point, and radiotracer; d: WT: *n*=14; HET: *n*=14

### [^18^F]CHDI-650 PET imaging reveals differences between genotypes

Striatal SUV TACs show stable uptake of [^18^F]CHDI-650 in the brain, with peak activity within 3 min p.i., with HET mice showing a slower wash-out than WT mice (both 9 months of age) (Fig. 3a). When performing kinetic modelling, several models were explored to fit the data. None of the tested compartmental models indicated a fully adequate fit for both WT and HET mice based on quantitative assessment and SE% exceeding 20%. Overall, when increasing the number of compartments in the model the model curve described the data even better as visually demonstrated in Fig. S5 and via the R^2^ value (coefficient of determination) in Table S6. The main issue, however, lay with the excessively high SE% of the estimates, especially with increased complexity of these models and even less reliable outcomes. Noteworthy, even though both WT and HET animals could not be described accurately by any compartmental model, the underlying issue slightly differed. For instance, WT data had a greater fit with 2TCM curves (R^2^: 0.97±0.01) than HET data (R^2^: 0.92±0.03) (Table S6). However, the SE% of the estimates, representing the essential challenge, were greater in WT than in HET animals (Table S6). 2TCM VascTrap was the only compartmental model with at least half of the animals considered valid for both genotypes. To identify a biological indication for this dichotomy, we investigated the presence of mHTT in the vasculature with immunostaining. However, as shown in Fig. S6, no co-localization of mHTT aggregates with the vascular tissue (i.e., lectin immunoreactivity) could be identified for any of the mHTT antibodies tested (mEM48, 2B4, and PHP1). As a fallback scenario, we explored the use of Logan graphical analysis as a simplified approach to obtain *V*_T(IDIF)_ as a quantifiable value. Given the suitability of Logan (Fig. 3b; 3c), we continued the assessment of [^18^F]CHDI-650 mindful of the caveat that comparative validation of Logan with a compartmental model was unavailable.

**Fig. 3.**
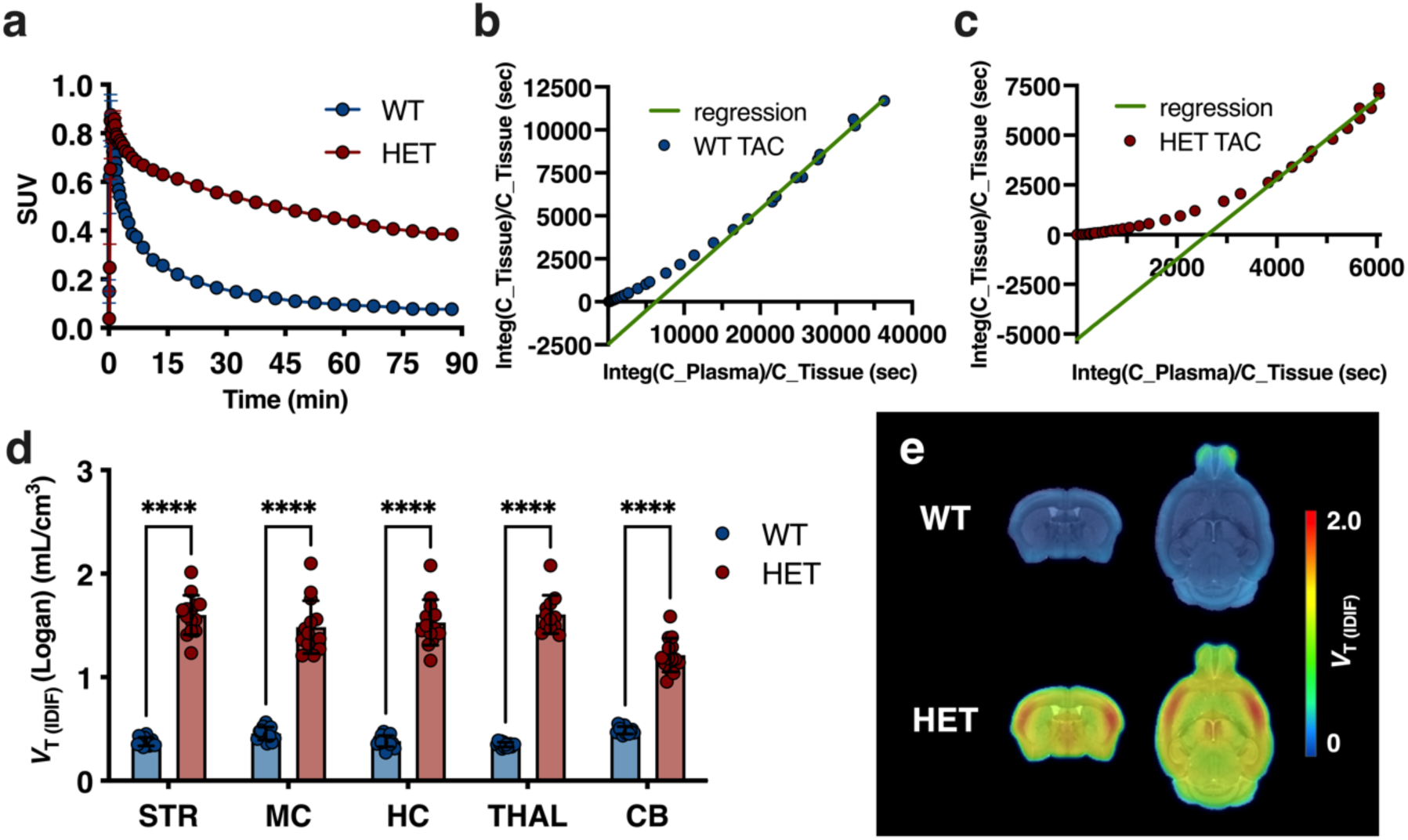
[^18^F]CHDI-650 mHTT quantification. a, Average striatal SUV TACs for both genotypes at 9 months of age. Exemplary WT (b) and HET (c) Logan fit. d, Significant differences between genotypes in *V*_T(IDIF)_ estimations. e, Group-averaged parametric maps for WT and HET zQ175DN mice overlaid onto MRI template in coronal and axial view. In a, data are represented as mean±SEM. \*\*\*\**p*<0.0001. a, d, and e: WT, *n*=12-14; HET, *n*=12-14. STR = Striatum; MC = Motor Cortex; HC = Hippocampus; THAL = Thalamus; CB = Cerebellum

To further assure proper mHTT aggregate imaging, parameters such as duration of the scan and injected mass of the radiotracer were investigated. The scans of [^18^F]CHDI-650 were acquired over a 90 min period; however, to increase the throughput rate with a single production and to lower the impact on animals, shortening the scan time is of interest. The *V*_T(IDIF)_ time stability analysis revealed that striatal Logan *V*_T(IDIF)_ values decreased to 82.39±2.91% in WT and 74.15±4.21% in HET for a shortened acquisition time of 60 min compared to 90 min reference scans (Fig. S7). Thus, 90 min was the recommended scan acquisition time for [^18^F]CHDI-650. With increasing injected masses, a decrease in *V*_T(IDIF)_ values was observed (Fig. S8); hence, the mass injection limit was set to 1 µg/kg for the inclusion of scans in all experiments reported here. By adhering to these recommendations, the two-way ANOVA (F_(1, 125)_=1739) performed with a Bonferroni post hoc test revealed a significant increase in signal in HET mice compared to their WT littermates in all tested regions (*p*<0.0001) (Fig. 3d) as also evidenced by the averaged parametric maps shown in Fig. 3e. Phenotypic *V*_T(IDIF)_ differences ranged from 149±12.1% in the cerebellum to 360±16.9% in the thalamus (Table 1).

**Table 1.** [^18^F]CHDI-650 *V*_T(IDIF)_ at 9 months of age. Data are given as mean±SD unless stated otherwise. WT, *n*=12-14; HET, *n*=12-14. Δ = difference between genotypes; SE = standard error

| | WT | HET | $p$ value | $\Delta\pm\text{SE}$ (%) |
| --- | --- | --- | --- | --- |
| **Striatum** | 0.38±0.04 | 1.60±0.19 | <0.0001 | 324±15.0 |
| **Motor Cortex** | 0.46±0.06 | 1.48±0.25 | <0.0001 | 225±12.6 |
| **Hippocampus** | 0.38±0.05 | 1.53±0.22 | <0.0001 | 299±14.8 |
| **Thalamus** | 0.35±0.02 | 1.61±0.18 | <0.0001 | 360±16.9 |
| **Cerebellum** | 0.49±0.04 | 1.21±0.16 | <0.0001 | 149±12.1 |

### [^18^F]CHDI-650 displays observable test-retest variability

The reproducibility of the imaging quantification was evaluated in a test-retest study in a different cohort than described in the previous paragraph. A comparison of the parametric maps for both genotypes at test and retest revealed no clear differences between the two scans of either WT or HET mice (Fig. 4a), with the regional quantifications of [^18^F]CHDI-650 shown in Table 2. Values for absolute test-retest variability in HET animals span from 13.1±12.0% in cerebellum to 17.7±11.0% in motor cortex. The absolute test-retest variability in WT animals ranged from 10.4±11.0% in cerebellum to 21.8±10.1% in striatum (Table 2), whereas the relative test-retest variability displayed an even distribution of *V*_T(IDIF)_ values for both genotypes between test and retest as displayed in a Bland-Altmann plot (Fig. 4b). In addition, values for individual animals varied from 28.2% to 33.4% for HET and from 16.5% to 37.6% for WT between test and retest, respectively. Finally, the ICC values for [^18^F]CHDI-650 were between 0.00 and 0.27 depending on the region (Table 2), which classified the test-retest performance of [^18^F]CHDI-650 as poor according to the reported classification [27].

**Fig. 4.**
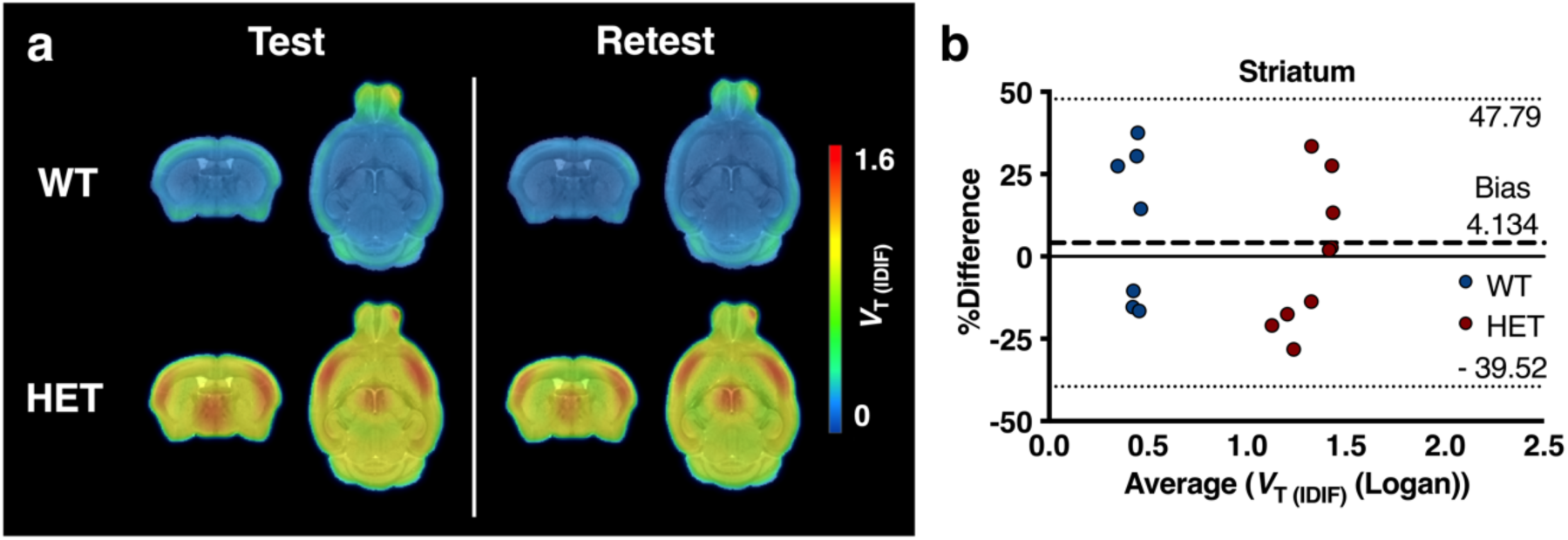
[^18^F]CHDI-650 test-retest variability analysis based on Logan quantification. a, Group-averaged parametric maps for WT and HET at test and retest at 9 months of age in coronal and axial view. b, Bland-Altmann-plot to display the level of agreement between test and retest in striatum. WT, *n*=4-7; HET, *n*=4-9

**Table 2.**
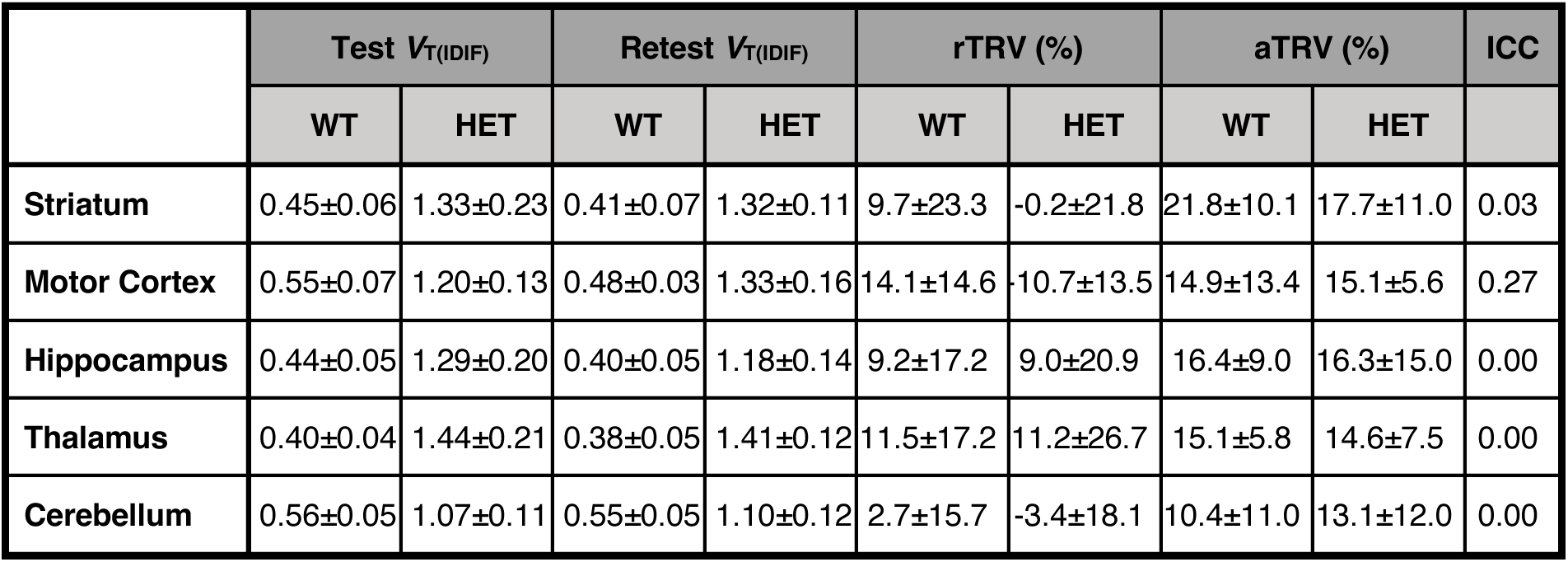
[^18^F]CHDI-650 test-retest *V*_T(IDIF)_ values and relative and absolute test-retest variability at 9 months of age. Test-retest variability is given as the average of the individual test-retest difference in percent and as the average of the absolute values of the individual test-retest difference in percent. The ICC is calculated for each region separately. WT, *n*=4-7; HET, *n*=4-9. rTRV = relative test-retest variability; aTRV = absolute test-retest variability; ICC = Intraclass correlation coefficient

| | Test $V_{T(\text{IDIF})}$ | | Retest $V_{T(\text{IDIF})}$ | | rTRV (%) | | aTRV (%) | | ICC |
| --- | --- | --- | --- | --- | --- | --- | --- | --- | --- |
|  | WT | HET | WT | HET | WT | HET | WT | HET |  |
| **Striatum** | 0.45±0.06 | 1.33±0.23 | 0.41±0.07 | 1.32±0.11 | 9.7±23.3 | -0.2±21.8 | 21.8±10.1 | 17.7±11.0 | 0.03 |
| **Motor Cortex** | 0.55±0.07 | 1.20±0.13 | 0.48±0.03 | 1.33±0.16 | 14.1±14.6 | -10.7±13.5 | 14.9±13.4 | 15.1±5.6 | 0.27 |
| **Hippocampus** | 0.44±0.05 | 1.29±0.20 | 0.40±0.05 | 1.18±0.14 | 9.2±17.2 | 9.0±20.9 | 16.4±9.0 | 16.3±15.0 | 0.00 |
| **Thalamus** | 0.40±0.04 | 1.44±0.21 | 0.38±0.05 | 1.41±0.12 | 11.5±17.2 | 11.2±26.7 | 15.1±5.8 | 14.6±7.5 | 0.00 |
| **Cerebellum** | 0.56±0.05 | 1.07±0.11 | 0.55±0.05 | 1.10±0.12 | 2.7±15.7 | -3.4±18.1 | 10.4±11.0 | 13.1±12.0 | 0.00 |

### [^18^F]CHDI-650 outperforms [^11^C]CHDI-180R in zQ175DN mice at 3 months of age

Finally, we performed a head-to-head comparison study with previously published [^11^C]CHDI-180R in zQ175DN mice at 3 months of age, an age that expresses significantly less aggregate load than the previously tested 9 months of age [13]. Average parametric maps for both [^18^F]CHDI-650 and [^11^C]CHDI-180R are shown in Fig. 5a and 5c. While both radioligands could demonstrate statistically significant differences between genotypes ([^18^F]CHDI-650: *F*_(1, 186)_=215.5, [^11^C]CHDI-180R: *F*_(1, 185)_=42.73), the magnitude of this difference was greater for [^18^F]CHDI-650 (for instance in thalamus: [^18^F]CHDI-650: 48.2±4.8%, *p*<0.0001; [^11^C]CHDI-180R: 11.2±2.2%, *p*<0.0001) (Fig. 5b; 5d and Table 3). These results were subsequently used to perform a power analysis (*α*=0.05, *β*=80%) to estimate the number of animals needed for statistical significance within future mHTT lowering therapy studies (Table 4). [^18^F]CHDI-650 displayed a higher effect size (measured as Cohen’s *d*) compared to [^11^C]CHDI-180R.

**Fig. 5.**
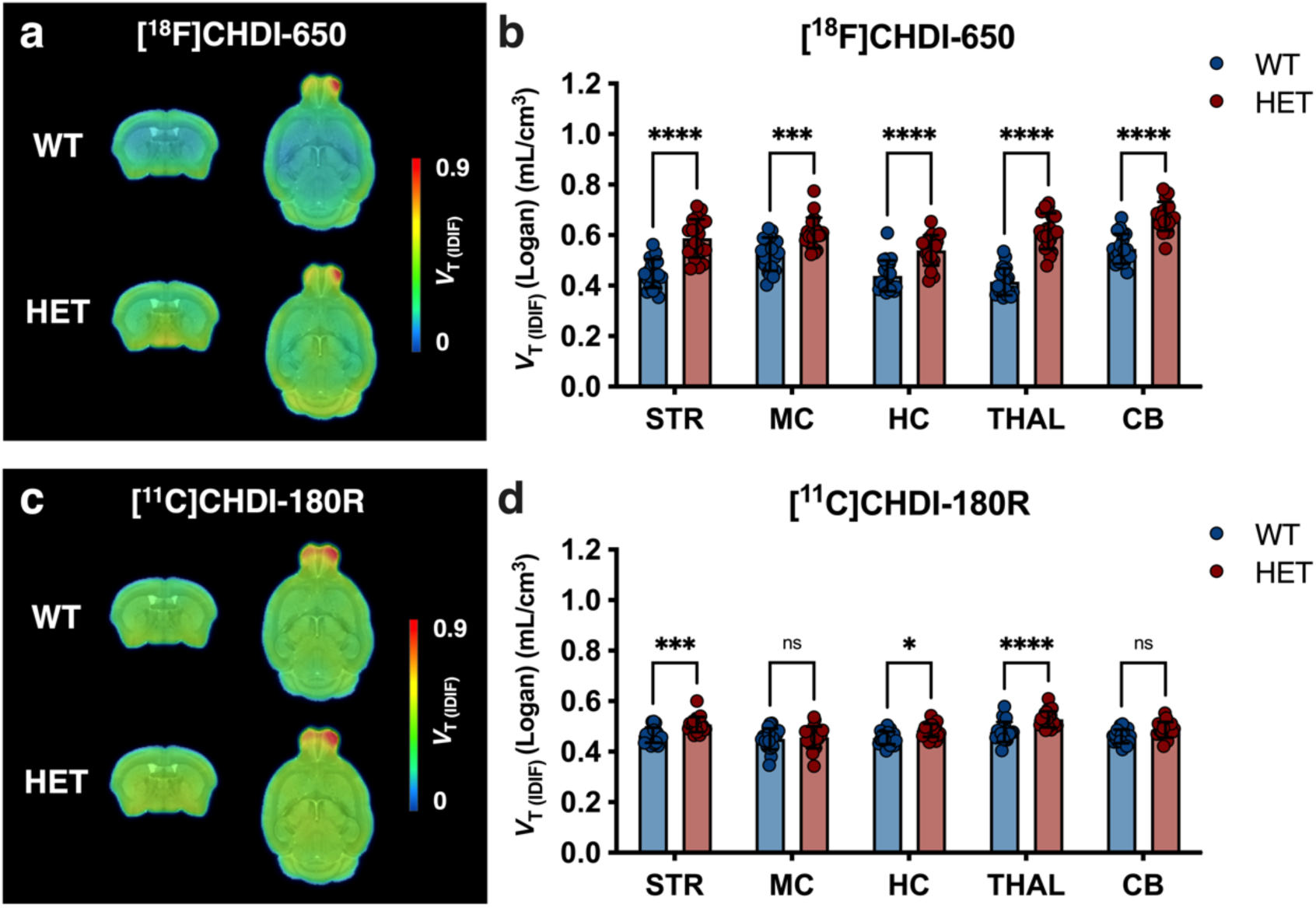
[^18^F]CHDI-650 and [^11^C]CHDI-180R comparison at 3 months of age. Group-averaged parametric maps for WT and HET mice displaying the *V*_T(IDIF)_ quantified with [^18^F]CHDI-650 (a) and [^11^C]CHDI-180R (c). Parametric maps are overlaid onto an MRI template and shown in coronal and axial views. Significant differences between genotypes in *V*_T(IDIF)_ estimations with [^18^F]CHDI-650 (b) and [^11^C]CHDI-180R (d). ns = not significant; \**p*<0.05; \*\*\**p*<0.001; \*\*\*\**p*<0.0001. [^18^F]CHDI-650: WT, *n*=19-20; HET, *n*=19-20. [^11^C]CHDI-180R: WT: *n*=19; HET: *n*=20. STR = Striatum; MC = Motor Cortex; HC = Hippocampus; THAL = Thalamus; CB = Cerebellum

**Table 3.** [^18^F]CHDI-650 and [^11^C]CHDI-180R *V*_T(IDIF)_ quantitative values at 3 months of age. Data is given as mean±SD unless stated otherwise. [^18^F]CHDI-650: WT, *n*=19-20; HET, *n*=19-20; [^11^C]CHDI-180R: WT: *n*=19; HET: *n*=20. Δ = difference between genotypes; SE = standard error

| | [$^{18}\text{F}$]CHDI-650 | | | | [$^{11}\text{C}$]CHDI-180R | | | |
| --- | --- | --- | --- | --- | --- | --- | --- | --- |
| | WT | HET | *p* value | $\Delta\pm\text{SE}$ (%) | WT | HET | *p* value | $\Delta\pm\text{SE}$ (%) |
| **Striatum** | 0.45±0.06 | 0.59±0.08 | <0.0001 | 31.0±4.4 | 0.47±0.03 | 0.51±0.03 | 0.0060 | 9.6±2.3 |
| **Motor Cortex** | 0.52±0.07 | 0.61±0.06 | 0.0002 | 16.1±3.8 | 0.45±0.04 | 0.46±0.04 | >0.9999 | 1.9±2.3 |
| **Hippocampus** | 0.44±0.06 | 0.54±0.06 | <0.0001 | 23.0±4.5 | 0.45±0.03 | 0.49±0.03 | 0.0269 | 7.2±2.3 |
| **Thalamus** | 0.42±0.05 | 0.62±0.07 | <0.0001 | 48.2±4.8 | 0.48±0.04 | 0.53±0.03 | <0.0001 | 11.2±2.2 |
| **Cerebellum** | 0.55±0.06 | 0.67±0.06 | <0.0001 | 23.6±3.7 | 0.46±0.03 | 0.49±0.03 | 0.0705 | 6.2±2.3 |

**Table 4.** [^18^F]CHDI-650 and [^11^C]CHDI-180R power analysis in striatum at 3 months of age. One-tailed power analysis with *α*=0.05 and *β*=80%. Animal numbers per group needed for significance between treated and non-treated HET group in a mHTT-lowering study. Percentages are the difference between these groups. 100% equals the difference between non-treated HET and WT groups. Δ = difference between genotypes; SE = standard error

|  |  |  | Phenotypic difference (%) |  |  |  |  |  |
| --- | --- | --- | --- | --- | --- | --- | --- | --- |
| | striatal $\Delta\pm\text{SE}$<br>(HET vs WT) | Cohen's *d* | 10% | 20% | 30% | 40% | 50% | 100% |
| **[$^{18}\text{F}$]CHDI-650** | 31.0±4.4% | 1.98 | $n=405$ | $n=102$ | $n=46$ | $n=26$ | $n=17$ | $n=5$ |
| **[$^{11}\text{C}$]CHDI-180R** | 9.6±2.3% | 1.33 | $n=697$ | $n=175$ | $n=77$ | $n=45$ | $n=29$ | $n=8$ |

## Discussion

The detailed *in vivo* characterisation of [^18^F]CHDI-650 for mHTT aggregates in the zQ175DN mouse model of HD is reported. We performed autoradiography, investigated the radiometabolite profile, and conducted an extensive characterisation for *in vivo* quantification in adult animals. Specifically, the radiometabolite profile was explored for both [^18^F]CHDI-650 and its non-deuterated version [^18^F]CHDI-961 to compare their *in vivo* metabolic stability. It is worth noting that [^18^F]CHDI-961 exhibited significant bone uptake, likely due to its propensity for oxidative defluorination, rendering it less favourable for brain imaging analysis [17]. For this reason, its deuterated form, [^18^F]CHDI-650 was profiled here in PET studies and supported by autoradiography.

The radiometabolite analysis of plasma revealed good metabolic stability for [^18^F]CHDI-650 following intravenous injection in zQ175DN mice. In addition, we verified the increased *in vivo* metabolic stability of [^18^F]CHDI-650 (30.4±5.1% at 45 min) compared to [^18^F]CHDI-961 (19.2±3.8% at 40 min) resulting, in part, from the addition of deuterium atoms that suppressed oxidative defluorination [17]. The plasma radiometabolite profiles of [^18^F]CHDI-650 in both genotypes generated at least two highly polar species of metabolites, which were not observed having a brain-penetrating behaviour, thus no impact on kinetic modelling is to be expected. Additionally, the intact [^18^F]CHDI-650 tracer showed a slower elimination from the brain of HET animals compared to WT controls, likely due to its binding to the mHTT aggregates, which is also seen with SUV TACs from the brain regions. The extracted brain signal demonstrated rapid penetration of [^18^F]CHDI-650 into the brain followed by a moderate wash-out of the tracer in HET zQ175DN mice and a faster wash-out reaching plateau around 60 min in WT littermates, indicating reversible binding in both genotypes. In contrast, [^11^C]CHDI-626 displayed rapid clearance from the brain of WT and HET mice [12], while [^11^C]CHDI-180R shows a long retention time within the HET brain [13].

The kinetic modelling was challenging as [^18^F]CHDI-650 could not be mathematically fitted to any compartmental models due to high SE% of the estimates or a divergence between the data and the model curve. Even though compartmental models with more parameters/compartments indicated a closer description of the data (i.e., higher R^2^ in HET animals), the increasing SE% of these additional parameters demonstrated the unreliability of these estimates. Significantly, WT zQ175DN mice presented excessively high SE%, suggesting that the lack of target for [^18^F]CHDI-650 in those animals may contribute to the inadequate compartmental model fits. Thus, [^18^F]CHDI-650 is reported without compartmental verification unlike [^11^C]CHDI-626 [12] and [^11^C]CHDI-180R [13]. Nonetheless, [^18^F]CHDI-650 can be adequately quantified with Logan analysis based on the 90 min acquisition. *V*_T(IDIF)_ quantification of [^18^F]CHDI-650 in 9-month-old animals suggests high binding in HET mice and low uptake in WT animals, reflecting the respective expression of mHTT aggregates in HET but not WT mice. This was also demonstrated using autoradiography. The potential of [^18^F]CHDI-650 to quantify mHTT aggregates is highlighted by the higher striatal *V*_T(IDIF)_ differences between genotypes (324±15%) compared to the two previously described ^11^C radiotracers for mHTT PET imaging, [^11^C]CHDI-626 (53.4±3.9%) [12] and [^11^C]CHDI-180R (63.1±2.8%) [13]. Recently, preclinical evaluation of another ^18^F radiotracer based on CHDI-180R has been reported [28]. This ^18^F analogue of CHDI-180R displayed rapid brain uptake but its *in vivo* specificity for mHTT aggregates remains unresolved; mHTT binding was reported through autoradiography analysis in *post-mortem* brain tissue of PwHD; however, there was evidence of white matter accumulation and non-specific binding [28]. On the other hand, Liu et al. showed through qualitative analysis of [^18^F]CHDI-650 a lack of non-specific binding in *post-mortem* brain tissue of PwHD and non-diseased controls [17]. Similarly, we could demonstrate this lack of binding of [^18^F]CHDI-650 in the absence of a target in WT zQ175DN at 9 months of age. Thus [^18^F]CHDI-650, appears to be a tractable radioligand for mHTT aggregate imaging.

The *in vivo* quantitative reliability of [^18^F]CHDI-650 was determined as poor through the test-retest variability analysis despite a low bias (4.3%) displayed in the Bland-Altmann plot. The ICC indicates how strongly units of the same group resemble each other, and in this study, ranged from 0.00 to 0.27 depending on the brain region, which is classified as poor [27]. Since HET and WT animals were affected, scan time duration and mHTT aggregate load could not be identified as the main factors to explain the reported test-retest variability. A less-than-optimal test-retest variability may negatively influence the estimated genotype effect. However, the high striatal *V*_T(IDIF)_ phenotypic difference of 324±15% at 9 months of age mitigates the potential effect on reduced statistical significance. Nevertheless, this may pose a challenge to its accuracy in monitoring longitudinal changes within the same animal.

As previously reported for [^11^C]CHDI-626 and [^11^C]CHDI-180R [12, 13], [^18^F]CHDI-650 also displayed significant differences in binding levels in HET animals compared to WT mice at 3 months of age. However, the variance and the standard deviation were reported higher for [^18^F]CHDI-650 than for [^11^C]CHDI-180R in the same animals, which was already hinted at during the test-retest variability evaluation at 9 months of age. Despite the high variability, [^18^F]CHDI-650 could discriminate between genotypes in all tested regions in contrast to [^11^C]CHDI-180R in the same animals. The results reported here concerning [^11^C]CHDI-180R not displaying significant genotype effects in all regions at 3 months of age are in line with previous findings [13]. The subsequent power analysis suggested that [^18^F]CHDI-650 could indicate significant therapeutic effects of at least a 40% decrease in mHTT aggregate levels with a considerably small number of animals per group at even 3 months of age ([^18^F]CHDI-650: *n*=26; [^11^C]CHDI-180R: *n*=45). Collectively, these findings suggest that [^18^F]CHDI-650 is a suitable ^18^F radioligand to image mHTT aggregates in living mice with limitations based on kinetic modelling and test-retest variability.

## Supplementary Information

Supplementary material to this article can be found at the end of this document.

## Declarations

## Acknowledgements

The authors acknowledge the contribution of the following CHDI staff: Drs Mette Skinbjerg, Robert Doot and Yuchuan Wang for helpful discussions; Dr Simon Noble for reviewing and providing constructive feedback; Brenda Lager for coordinating animal supply and Maya Bader for project management. The authors thank Philippe Joye, Caroline Berghmans, Romy Raeymakers, and Eleni Van der Hallen of the Molecular Imaging Center Antwerp (MICA) for their valuable assistance.

## Authors contributions

Conceptualisation: Daniele Bertoglio, Jeroen Verhaeghe, Ignacio Munoz-Sanjuan, Celia Dominguez, Vinod Khetarpal, Jonathan Bard, Longbin Liu, Steven Staelens; Methodology: Daniele Bertoglio, Jeroen Verhaeghe, Alan Miranda, Steven Staelens; Data acquisition: Franziska Zajicek, Stef De Lombaerde, Annemie Van Eetveldt; Formal analysis and investigation: Franziska Zajicek, Annemie Van Eetveldt; Data interpretation: Franziska Zajicek, Jeroen Verhaeghe, Stef De Lombaerde, Ignacio Munoz-Sanjuan, Vinod Khetarpal, Jonathan Bard, Longbin Liu, Daniele Bertoglio, Steven Staelens; Writing – original draft preparation: Franziska Zajicek, Daniele Bertoglio, Steven Staelens; Writing – review and editing: Franziska Zajicek, Jeroen Verhaeghe, Stef De Lombaerde, Alan Miranda, Ignacio Munoz-Sanjuan, Vinod Khetarpal, Jonathan Bard, Longbin Liu, Daniele Bertoglio, Steven Staelens.

## Ethics approval

Experiments followed the European Committee (decree 2010/63/CEE) and were approved by the Ethical Committee for Animal Testing (ECD 2019-39) at the University of Antwerp (Belgium).

## Supplemental information

### Immunohistochemistry

Immunohistochemistry was performed on PFA-perfused frozen tissue of 9-month-old WT and HET zQ175DN animals in order to explore the presence of mHTT aggregates in the proximity of the vasculature within the brain. Three different antibodies for mHTT were used for the investigation of endothelial binding, namely PHP1 (CHDI, CH01951), EM48 (Merck, MAB5374), and 2B4 (Merck, MAB5492), which bind to different parts and inclusions of mHTT and therefore aided in displaying a more complete picture of the co-localization of the vasculature and mHTT.

Staining with Lectin and PHP1 was performed as follows: sections were acclimatised at room temperature (RT) for 5 min and subsequently rinsed with 1x PBS for another 5 min. Slides were then washed three consecutive times with 1x PBS for 5 min. Then, they were incubated in 1% formic acid for 10 min to decalcify the sections followed by three washing steps, twice with PBS + 0.25% Triton X-100 for 5 min and once with PBS for 5 min. Excess fluid was then disregarded and slides were incubated with 5% Normal Donkey Serum (NDS) + 0.5% Triton X-100 in 1x PBS for 30 min. After letting excess fluid drop off, a 1 h blocking step to avoid non-specific binding to endogenous mouse IgG using Donkey anti-mouse Fab Fragment IgG (10-20 µg/ml in PBS) was performed followed by overnight incubation with the primary antibody mouse anti-mouse mHTT PHP1 1:3000 (CHDI, CH01951) in 1% NDS + 0.1% Triton X-100 in 1x PBS at RT. The following day, sections were washed three times with 1x PBS for 5 min prior to incubation with the secondary antibody donkey anti-mouse AF555 (1:1000) for 1 h. Sections were then washed another three times with 1x PBS for 5 min before incubation with Lectin Dylight488 1:25 (Vector laboratories, DL-1174) in PBS for 2 h. After a final washing step with 1x PBS, DAPI was added, and the sections were mounted with prolong gold antifade and coverslipped.

Staining with Lectin and EM48 or 2B4 was performed as follows: Sections were acclimatised at RT for 5 min and subsequently rinsed with 1x PBS for another 5 min. Slides were then put in a container with citrate buffer and the container in turn in a 60°C water bath. Slides were incubated for 20 min as soon as the water bath reached the final temperature of 80°C. A subsequent cool-down step in the citrate buffer for 20 min, as well as three washing cycles with 1x PBS for 5 min, preceded the non-specific binding blockage of endogenous proteins with 0.5% Triton X-100 + 10% NDS in PBS for 30 min. Endogenous mouse IgG blocking was performed with Donkey anti-mouse Fab Fragment IgG (10-20 µg/ml) in PBS for 1 h. Slides were then washed again prior to the overnight incubation at RT with the primary antibody anti-mouse EM48 (1:200) (Merck, MAB5374, IgG) in PBS + 1% NDS + 1% BSA + 0.1% Triton X-100 or anti-mouse 2B4 (1:500) (Merck, MAB5492, IgG1) in PBS + 1% NDS + 0.1% Triton X-100.

The next day slides were incubated with the secondary antibody Donkey anti-mouse AF555 (1:1000) in PBS (+ 0.2% BSA for EM48) for 1 h after washing three times with 1x PBS for 5 min. Sections were then washed another three times with 1x PBS for 5 min before incubation with Lectin Dylight488 1:25 (Vector laboratories, DL-1174) in PBS for 2 h. After a final washing step with 1x PBS, DAPI was added, and the sections were mounted with prolong gold antifade and coverslipped.

## Supplemental Figures

**Fig. S1.**
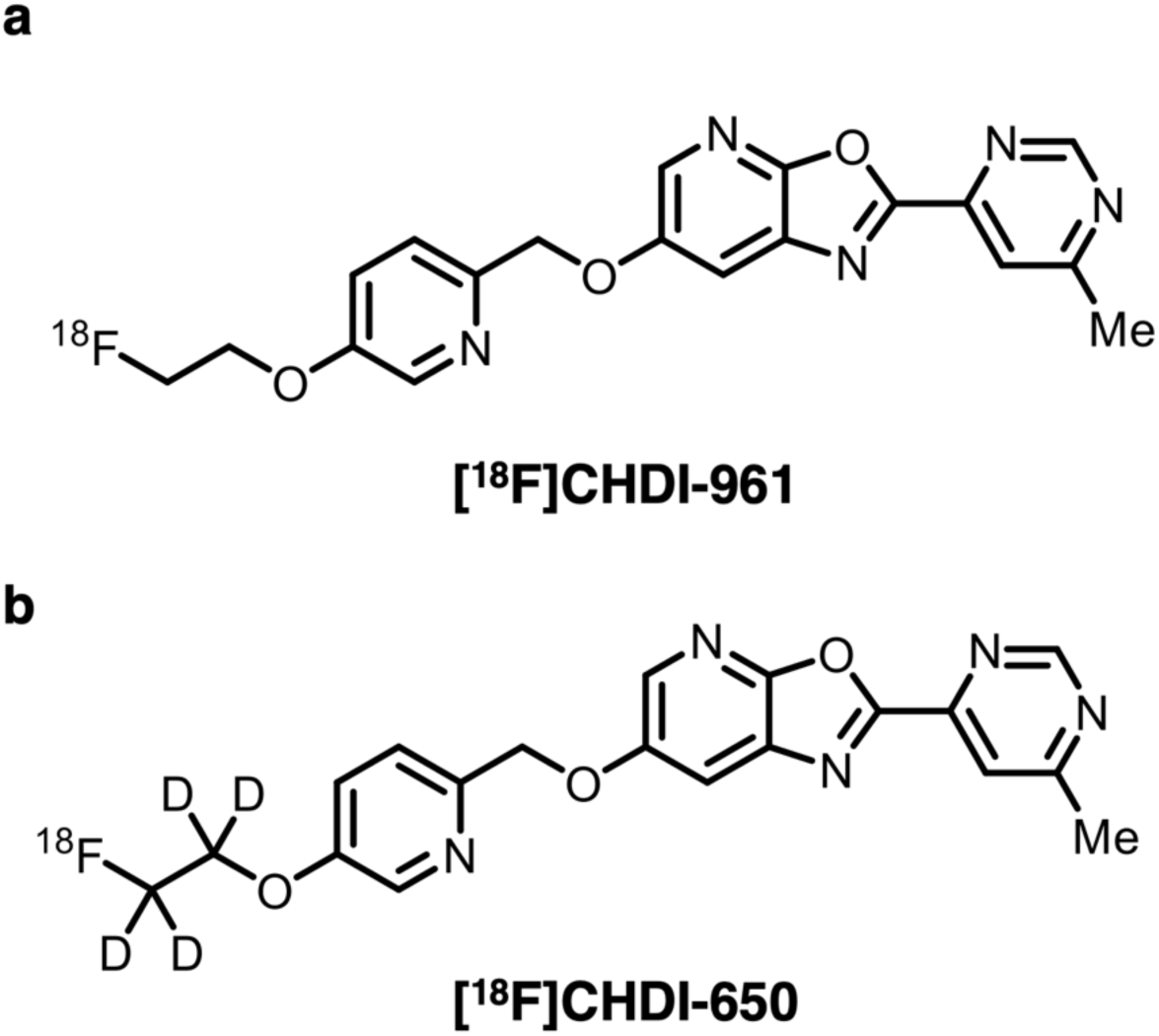
Molecular structure of [^18^F]CHDI-961 and [^18^F]CHDI-650. [^18^F]CHDI-961 (a) resulted in [^18^F]CHDI-650 (b) through the addition of Deuterium at the ^18^F site

**Fig. S2.**
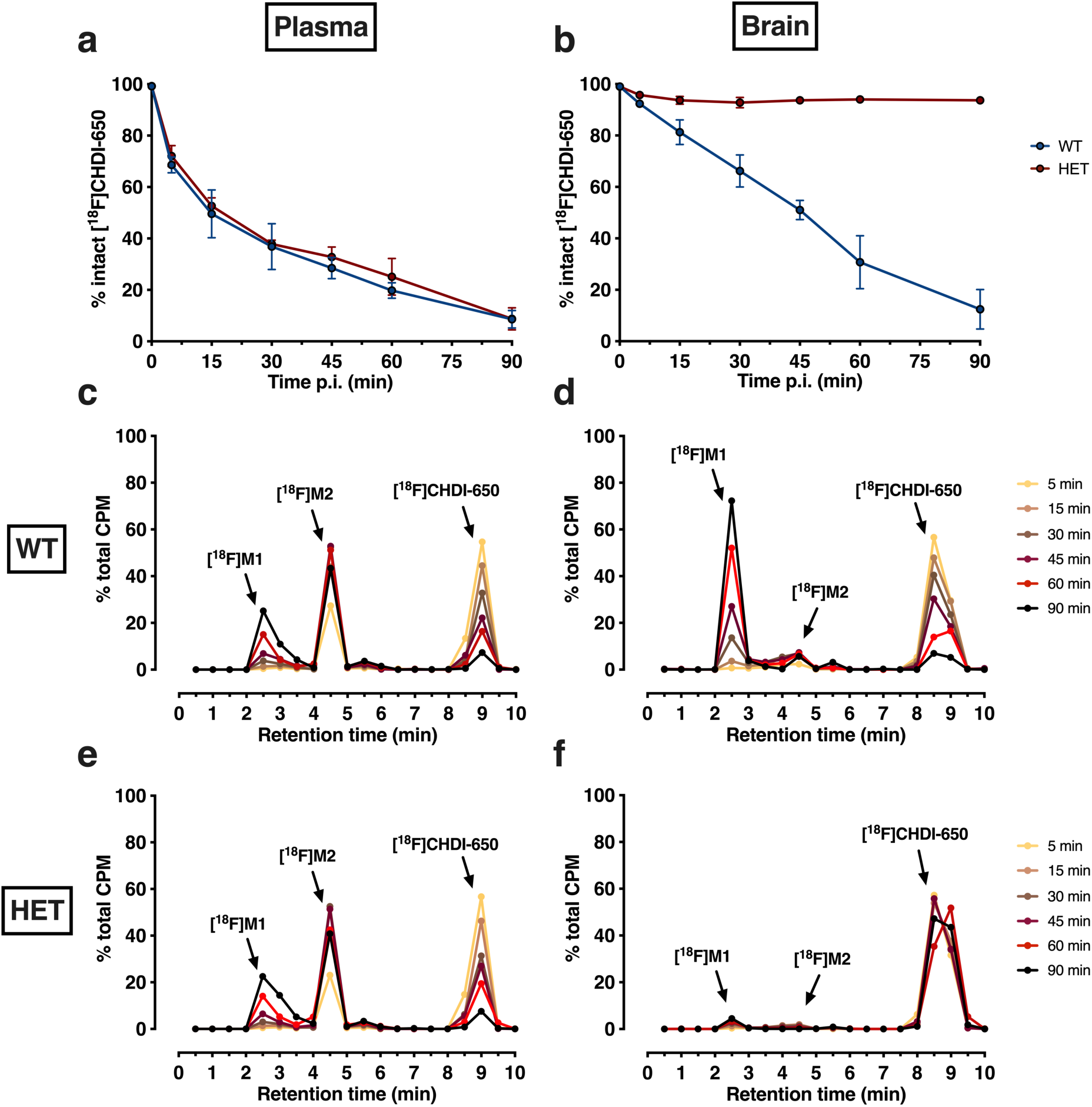
[^18^F]CHDI-650 plasma and brain radiometabolite profile and decay-corrected reconstructed radiochromatograms in WT and HET zQ175DN mice for plasma and brain. The intact radioligand in percentage for plasma (a) and brain (b) displayed a similar decrease in plasma between genotypes and the stability in HET brains compared to the decrease in WT brains over time. c-f; decay-corrected reconstructed radiochromatograms for all time points in plasma (c and e) and brain (d and f) for WT (c and d) and HET mice (e and f). *n*=3-7 per genotype, time point, and radiotracer. [^18^F]M1 = radiometabolite peak 1; [^18^F]M2 = radiometabolite peak 2

**Fig. S3.**
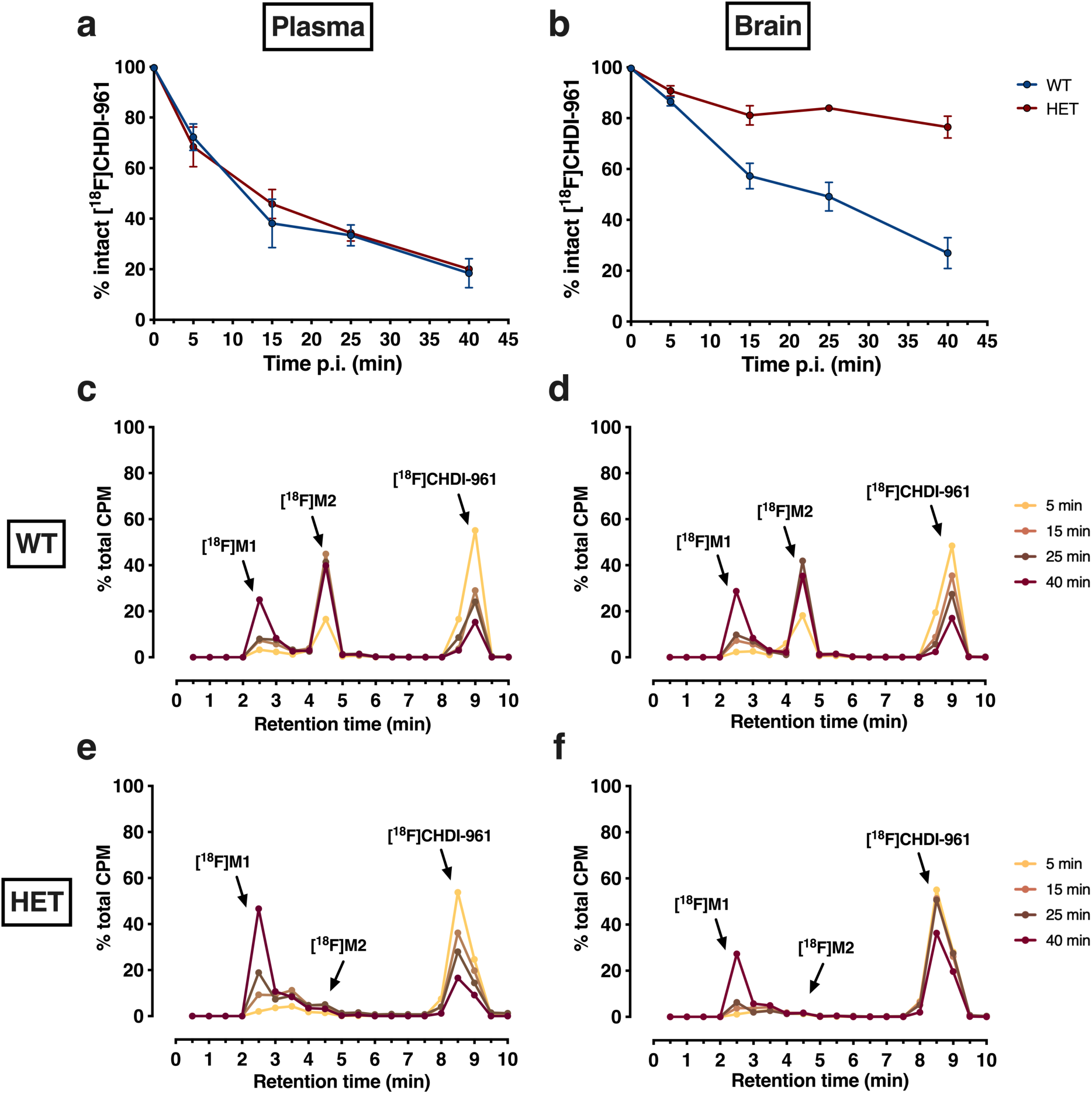
[^18^F]CHDI-961 plasma and brain radiometabolite profile and decay-corrected reconstructed radiochromatograms in WT and HET zQ175DN mice for plasma and brain. The intact radioligand in percentage for plasma (a) and brain (b) displayed a similar decrease in plasma between genotypes and the stability in HET brains compared to the decrease in WT brains over time. c-f; decay-corrected reconstructed radiochromatograms for all time points in plasma (c and e) and brain (d and f) for WT (c and d) and HET mice (e and f). *n*=2-6 per genotype, time point, and radiotracer. [^18^F]M1 = radiometabolite peak 1; [^18^F]M2 = radiometabolite peak 2

**Fig. S4.**
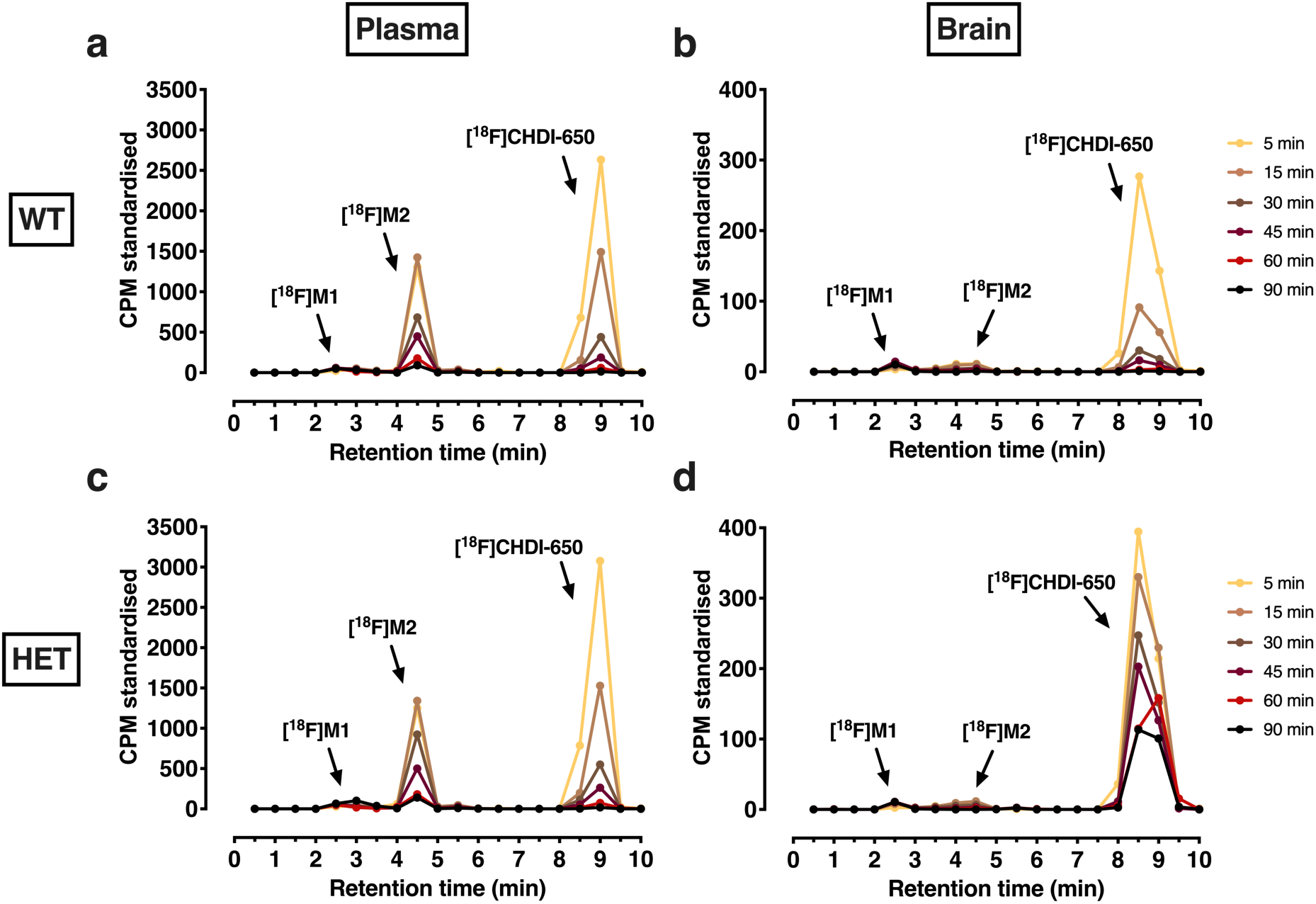
[^18^F]CHDI-650 plasma and brain decay-corrected reconstructed standardised radiochromatograms in WT (a and b) and HET (c and d) zQ175DN mice for plasma (a and c) and brain (b and d). *n*=3-7 per genotype, time point, and radiotracer. [^18^F]M1 = radiometabolite peak 1; [^18^F]M2 = radiometabolite peak 2

**Fig. S5.**
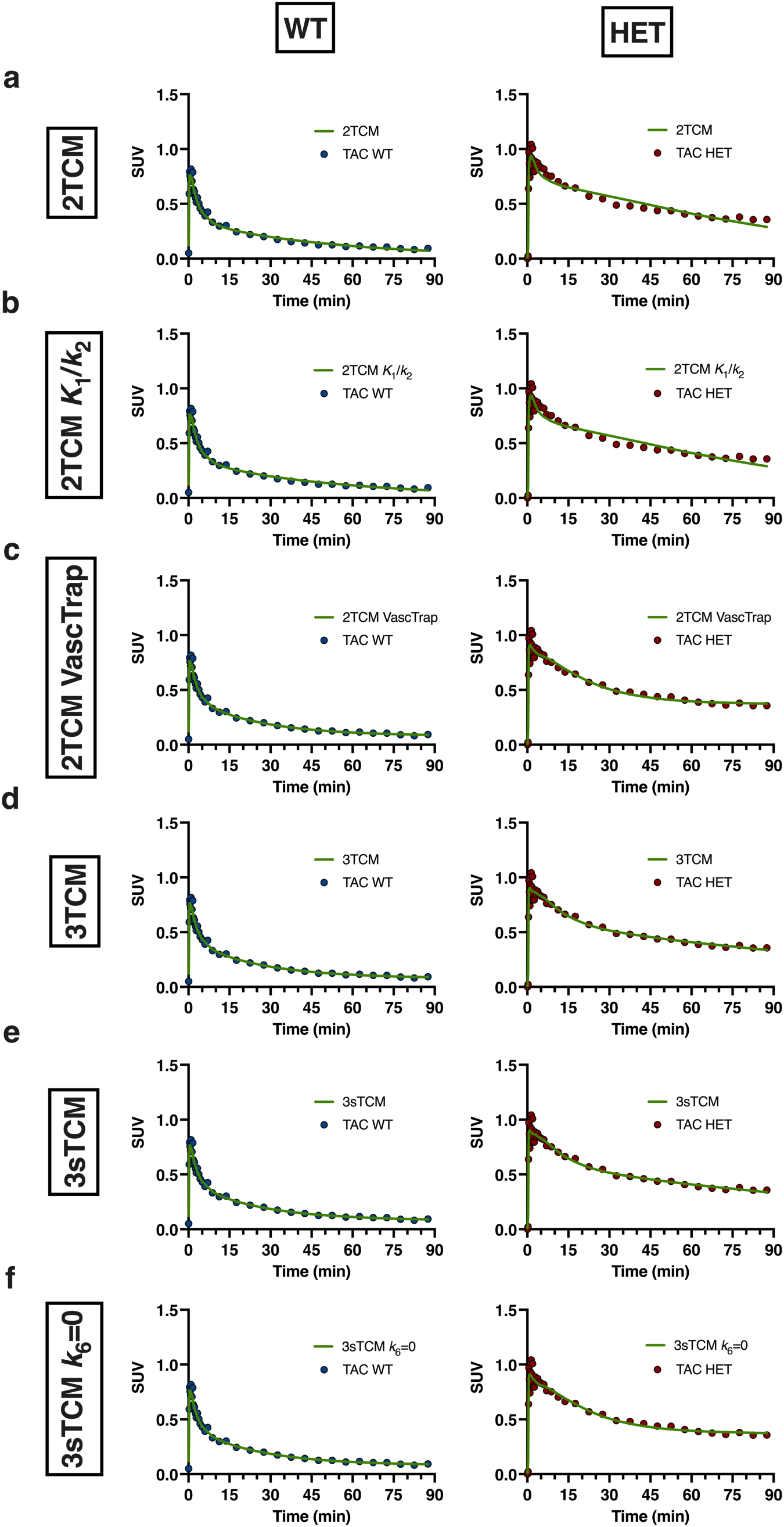
Kinetic model fits in both genotypes. Exemplary model fits of striatal SUV TACs of a WT and a HET zQ175DN mouse with 2TCM (a), 2TCM *K*_1_/*k*_2_ (b), 2TCM VascTrap (c), 3TCM (d), 3sTCM (e), 3sTCM *k*_6_=0 (f)

**Fig. S6.**
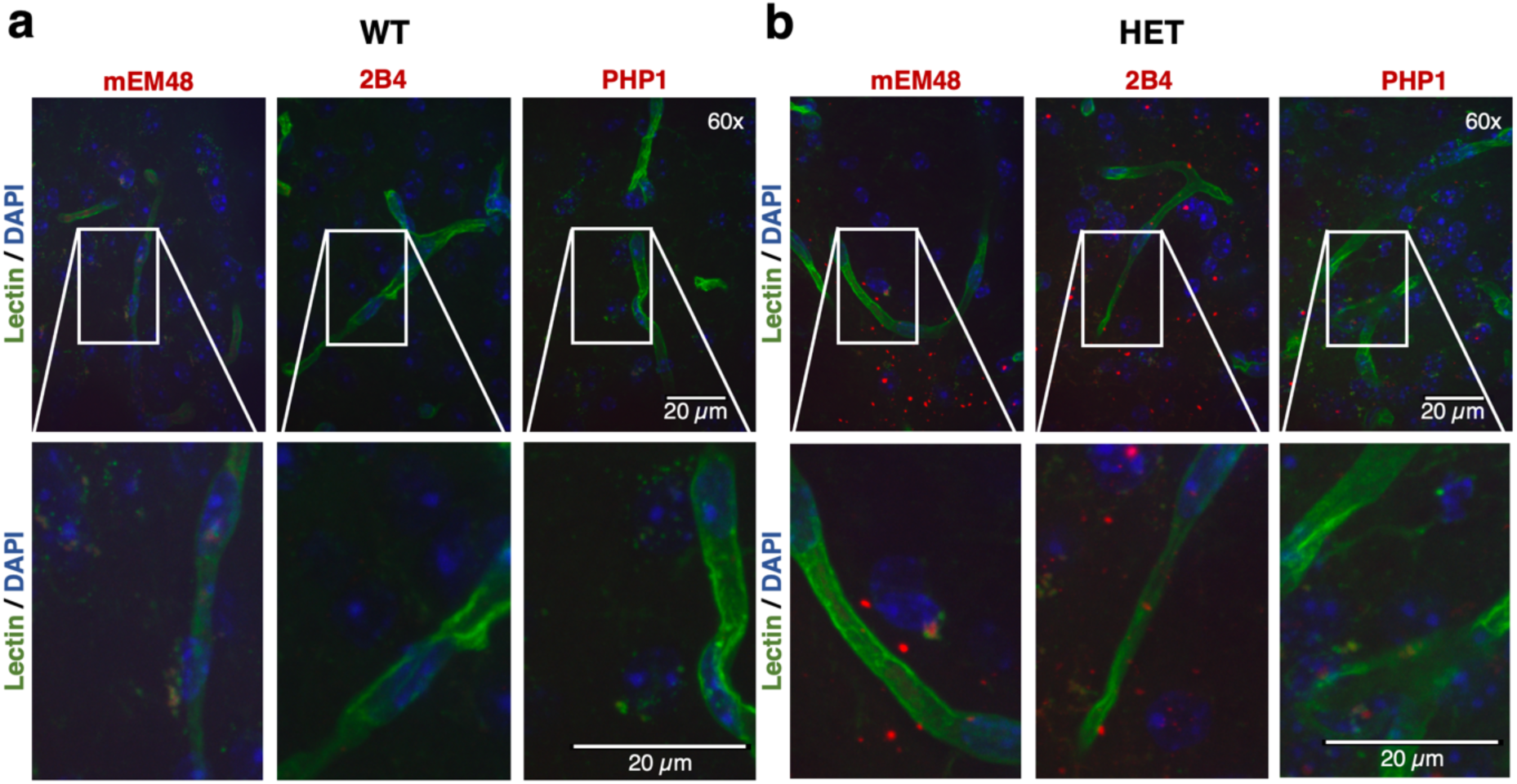
mHTT aggregates do not co-localise with vasculature tissue in brain of zQ175DN mice at 9 months of age. Exemplary images of striatal tissue of WT (a) and HET (b) animals stained with Lectin (green) and either mEM48, 2B4 or PHP1 (red). DAPI staining (blue) to visualise nuclei. Images are displayed with 60x magnification with relevant areas further magnified. Scale indicates 20µm

**Fig. S7.**
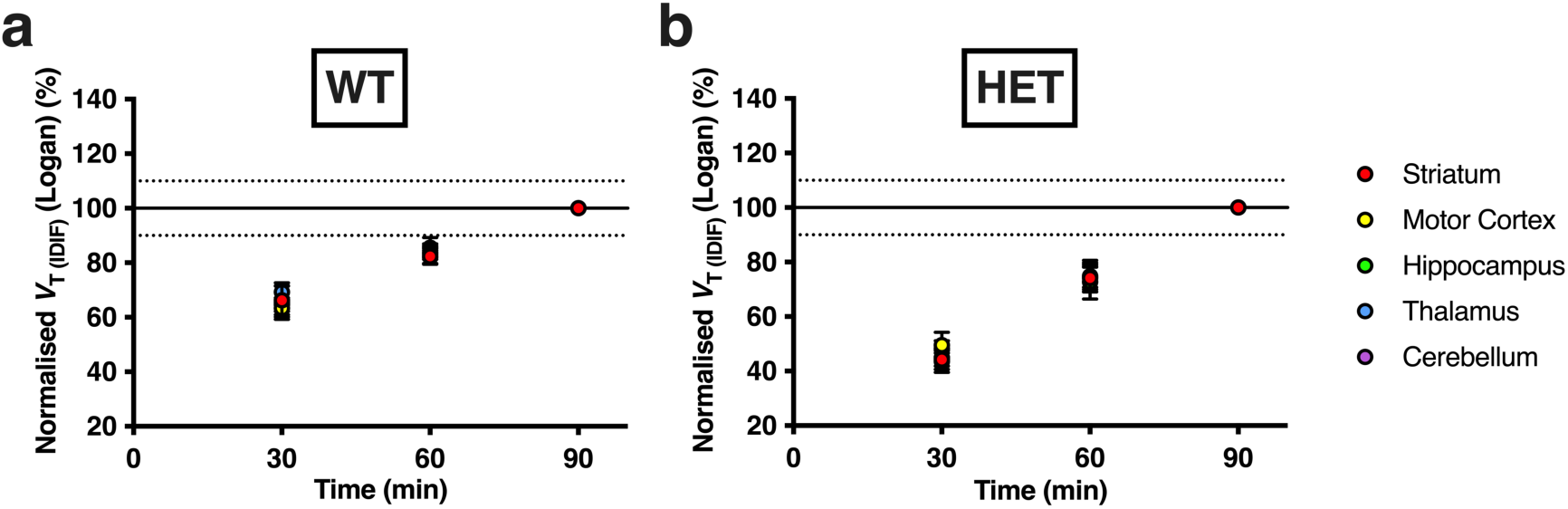
*V*_T(IDIF)_ time stability in WT and HET zQ175DN mice. The *V*_T(IDIF)_ values estimated with Logan in WT (a) and HET (b) for 30 and 60 min are normalised to 90 min scan acquisition time. Data are represented as mean±SD in percentage. WT: *n*=12-14; HET: *n*=12-14

**Fig. S8.**
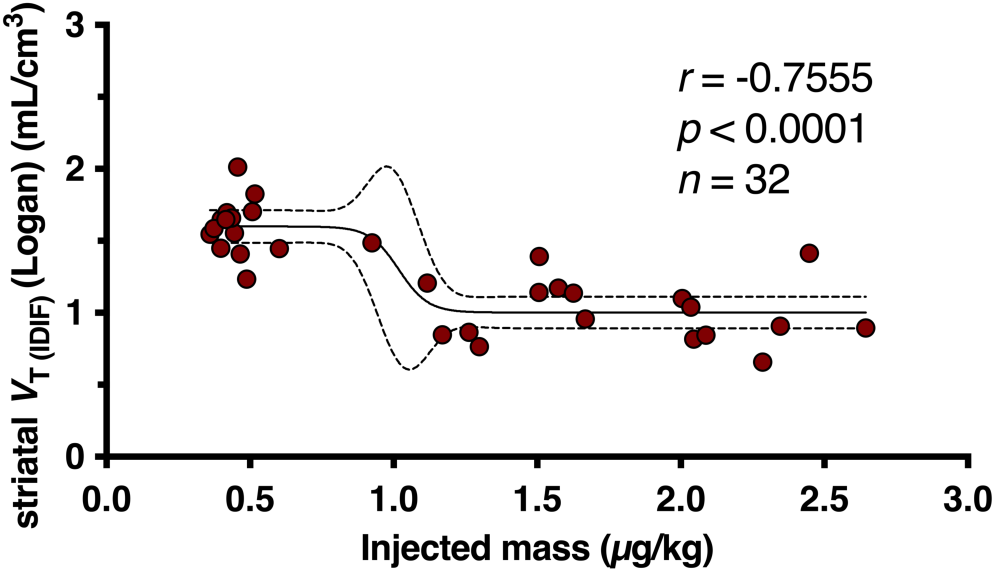
Mass dose effect in HET zQ175DN mice. Negative correlation of injected masses (µg/kg) and striatal Logan *V*_T(IDIF)_ values. The interpolation estimates the influence of higher injected masses on the *V*_T(IDIF)_ value. Data is represented as individual points. *n*=32

## Supplemental Tables

**Table S1.** Animal and dosing parameters during the PET imaging study in WT and HET zQ175DN mice for the validation at 9 months of age. Values are given as mean±SD. Unpaired t-test for comparison of genotypes. EOS=end of synthesis

|  | WT (*n* = 14) | HET (*n* = 14) | *p* value |
| --- | --- | --- | --- |
| Injected activity (MBq) | 7.0±1.9 | 6.9±3.1 | 0.9166 |
| Molar activity (GBq/μmol) at EOS | 236.1±70.9 | 221.5±69.5 | 0.5856 |
| Weight (g) | 34.6±3.3 | 31.1±3.0 | 0.0067 |
| Age (d) | 277.4±7.5 | 278.0±9.2 | 0.8405 |
| Injected mass (μg/kg) | 0.47±0.07 | 0.45±0.06 | 0.4139 |

**Table S2.** Animal and dosing parameters during the PET imaging study in WT and HET zQ175DN mice for the test-retest study at 9 months of age. Values are given as mean±SD. Paired t-test for comparison of test and retest. EOS=end of synthesis

|  | Test | Retest |  | Test | Retest |  |
| --- | --- | --- | --- | --- | --- | --- |
|  | WT (*n* = 7) |  | *p* value | HET (*n* = 9) |  | *p* value |
| Injected activity (MBq) | 6.5±2.2 | 6.9±1.9 | 0.7817 | 6.9±1.5 | 6.7±2.4 | 0.8249 |
| Molar activity (GBq/μmol) at EOS | 187.0±71.0 | 207.7±91.0 | 0.6491 | 230.0±79.1 | 233.8±92.3 | 0.9921 |
| Weight (g) | 34.3±1.5 | 33.3±0.9 | 0.0089 | 30.7±2.5 | 29.8±2.4 | 0.0075 |
| Age (d) | 254.4±1.0 | 261.1±3.4 | 0.0016 | 252.4±5.8 | 260.6±10.6 | 0.0013 |
| Injected mass (μg/kg) | 0.55±0.08 | 0.48±0.06 | 0.0893 | 0.53±0.08 | 0.52±0.09 | 0.8974 |

**Table S3.** Animal and dosing parameters during the PET imaging study in WT and HET zQ175DN mice for the 3 months of age head-to-head comparison study with [^18^F]CHDI-650 (A) and [^11^C]CHDI-180R (B). Values are given as mean±SD. Unpaired t-test for comparison of genotypes for each radiotracer. EOS=end of synthesis

|  | **[<sup>18</sup>F]CHDI-650** |  |  | **[<sup>11</sup>C]CHDI-180R** |  |  |
| --- | --- | --- | --- | --- | --- | --- |
|  | **WT (n = 20)** | **HET (n = 20)** | **p value** | **WT (n = 19)** | **HET (n = 20)** | **p value** |
| **Injected activity (MBq)** | 6.0±1.6 | 6.7±2.2 | 0.2127 | 2.2±0.3 | 2.2±0.3 | 0.9020 |
| **Molar activity (GBq/μmol) at EOS** | 241.1±60.7 | 257.2±61.6 | 0.4116 | 59.5±9.9 | 59.6±9.4 | 0.9593 |
| **Weight (g)** | 26.5±1.2 | 26.8±1.1 | 0.4321 | 27.2±1.1 | 26.5±1.3 | 0.0787 |
| **Age (d)** | 104.4±5.6 | 106.0±7.0 | 0.4286 | 116.2±2.9 | 115.9±3.1 | 0.7550 |
| **Injected mass (μg/kg)** | 0.54±0.07 | 0.52±0.06 | 0.4036 | 0.84±0.19 | 0.87±0.21 | 0.5568 |

**Table S4.** Animal numbers for the radiometabolite profile analysis. Time indicates the blood and brain extraction p.i.; Ctrl = control experiments

|  | <sup>18</sup>F]CHDI-961 |  |  |  | <sup>18</sup>F]CHDI-650 |  |  |  |
| --- | --- | --- | --- | --- | --- | --- | --- | --- |
|  | Plasma |  | Brain |  | Plasma |  | Brain |  |
| Time (min) | WT | HET | WT | HET | WT | HET | WT | HET |
| Ctrl | 2 | 2 | 2 | 2 | 3 | 3 | 3 | 3 |
| 5 | 3 | 3 | 3 | 3 | 3 | 3 | 3 | 3 |
| 15 | 5 | 6 | 5 | 6 | 4 | 4 | 4 | 4 |
| 25 | 3 | 4 | 3 | 4 |  |  |  |  |
| 30 |  |  |  |  | 6 | 4 | 7 | 4 |
| 40 | 3 | 3 | 3 | 3 |  |  |  |  |
| 45 |  |  |  |  | 5 | 4 | 5 | 4 |
| 60 |  |  |  |  | 5 | 5 | 5 | 5 |
| 90 |  |  |  |  | 4 | 4 | 3 | 4 |

**Table S5.** Animal numbers for dynamic PET imaging. ° = excluded due to spill-in; * = excluded due to high SE%. T-RT = test-retest

|  | 9M |  |  |  | 3M |  |  |  |
| --- | --- | --- | --- | --- | --- | --- | --- | --- |
|  | Quantification |  | T-RT analysis |  | <sup>18</sup>F]CHDI-650 |  | <sup>11</sup>C]CHDI-180R |  |
|  | WT<br>(n = 14) | HET<br>(n = 14) | WT<br>(n = 7) | HET<br>(n = 9) | WT<br>(n = 20) | HET<br>(n = 20) | WT<br>(n = 19) | HET<br>(n = 20) |
| Striatum | 14 | 14 | 7 | 9 | 20 | 20 | 19 | 20 |
| Motor Cortex | 13 (1°) | 14 | 4 (3°) | 4 (3°, 2*) | 20 | 19 (1*) | 19 | 20 |
| Hippocampus | 14 | 14 | 7 | 8 (1*) | 20 | 20 | 19 | 20 |
| Thalamus | 14 | 12 (2*) | 7 | 8 (1*) | 20 | 19 (1*) | 19 | 20 |
| Cerebellum | 12 (2°) | 14 | 5 (2°) | 8 (1°) | 19 (1°) | 19 (1°) | 19 | 20 |

**Table S6.**
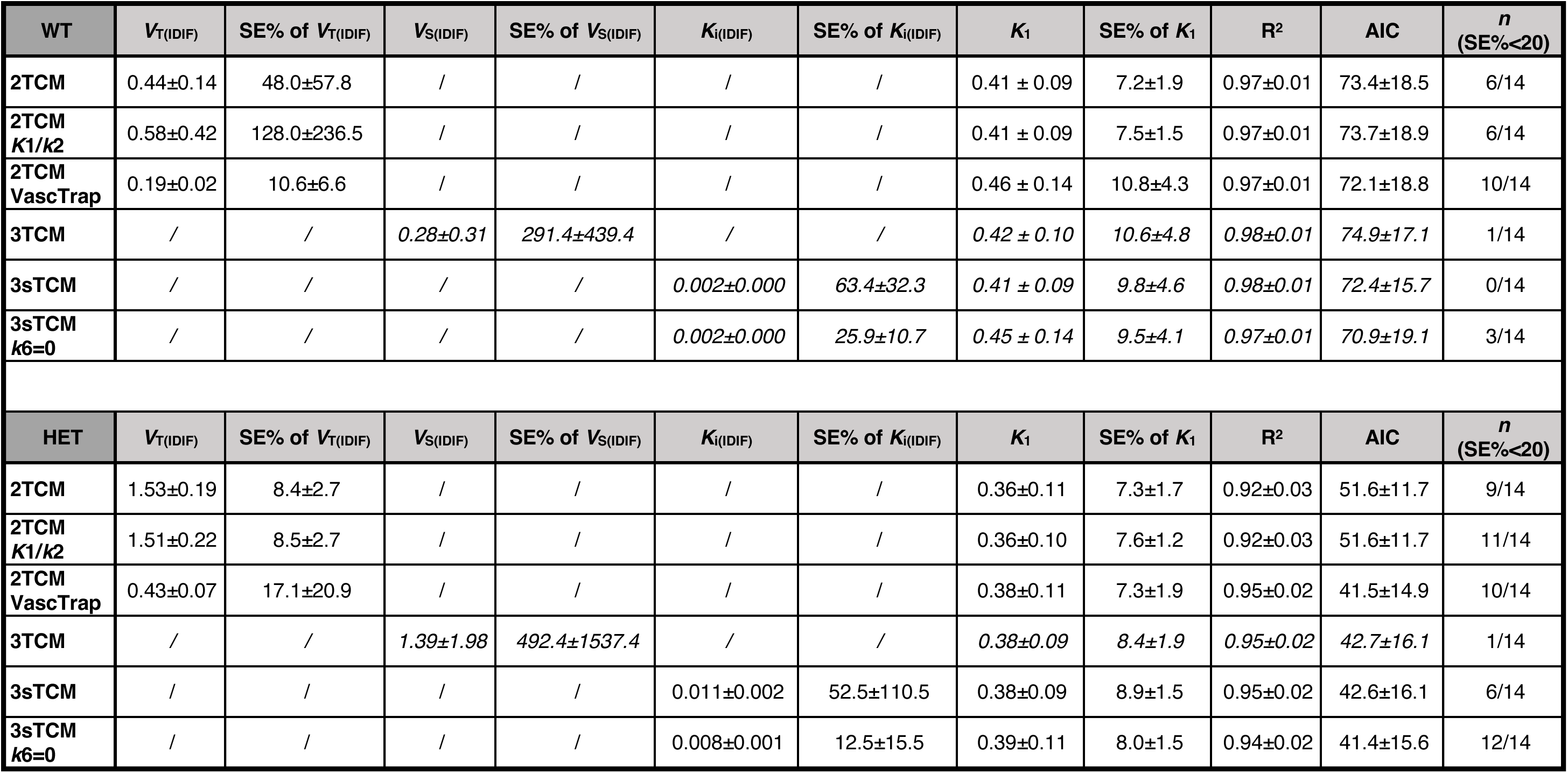
Kinetic model parameters and their standard errors for all investigated compartmental models in Striatum. Values are given as mean±SD or as valid animal number/total animal number. Exclusion of values higher than 10^5^ or lower than 10^-5^. n = valid animal number after exclusion of values due to SE% above 20 for given parameters and visual assessment. SE% = standard error percentage; AIC = Akaike information criterion. Unreliable values are indicated in italics

## References

1. Saudou F, Humbert S. The Biology of Huntingtin. Neuron. 2016;89:910–26. doi:10.1016/j.neuron.2016.02.003.

2. The-Huntington’s-Disease-Collaborative-Research-Group. A novel gene containing a trinucleotide repeat that is expanded and unstable on Huntington’s disease chromosomes. Cell. 1993;72:971–83. doi:10.1016/0092-8674(93)90585-e.

3. Waldvogel HJ, Kim EH, Tippett LJ, Vonsattel JP, Faull RL. The Neuropathology of Huntington’s Disease. Curr Top Behav Neurosci. 2015;22:33–80. doi:10.1007/7854_2014_354.

4. Tabrizi SJ, Ghosh R, Leavitt BR. Huntingtin Lowering Strategies for Disease Modification in Huntington’s Disease. Neuron. 2019;101:801–19. doi:10.1016/j.neuron.2019.01.039.

5. Wild EJ, Tabrizi SJ. Therapies targeting DNA and RNA in Huntington’s disease. The Lancet Neurology. 2017;16:837–47. doi:10.1016/s1474-4422(17)30280-6.

6. Hensman Moss DJ, Robertson N, Farmer R, Scahill RI, Haider S, Tessari MA, et al. Quantification of huntingtin protein species in Huntington’s disease patient leukocytes using optimised electrochemiluminescence immunoassays. PLoS One. 2017;12:e0189891. doi:10.1371/journal.pone.0189891.

7. Macdonald D, Tessari MA, Boogaard I, Smith M, Pulli K, Szynol A, et al. Quantification assays for total and polyglutamine-expanded huntingtin proteins. PLoS One. 2014;9:e96854. doi:10.1371/journal.pone.0096854.

8. Reindl W, Baldo B, Schulz J, Janack I, Lindner I, Kleinschmidt M, et al. Meso scale discovery-based assays for the detection of aggregated huntingtin. PLoS One. 2019;14:e0213521. doi:10.1371/journal.pone.0213521.

9. Liu L, Prime ME, Lee MR, Khetarpal V, Brown CJ, Johnson PD, et al. Imaging Mutant Huntingtin Aggregates: Development of a Potential PET Ligand. J Med Chem. 2020;63:8608–33. doi:10.1021/acs.jmedchem.0c00955.

10. Liu L, Johnson PD, Prime ME, Khetarpal V, Lee MR, Brown CJ, et al. [(11)C]CHDI-626, a PET Tracer Candidate for Imaging Mutant Huntingtin Aggregates with Reduced Binding to AD Pathological Proteins. J Med Chem. 2021;64:12003–21. doi:10.1021/acs.jmedchem.1c00667.

11. Herrmann F, Hessmann M, Schaertl S, Berg-Rosseburg K, Brown CJ, Bursow G, et al. Pharmacological characterization of mutant huntingtin aggregate-directed PET imaging tracer candidates. Sci Rep. 2021;11:17977. doi:10.1038/s41598-021-97334-z.

12. Bertoglio D, Verhaeghe J, Miranda A, Wyffels L, Stroobants S, Mrzljak L, et al. Longitudinal preclinical evaluation of the novel radioligand [11C]CHDI-626 for PET imaging of mutant huntingtin aggregates in Huntington’s disease. Eur J Nucl Med Mol Imaging. 2022;49:1166–75. doi:10.1007/s00259-021-05578-8.

13. Bertoglio D, Bard J, Hessmann M, Liu L, Gartner A, De Lombaerde S, et al. Development of a ligand for in vivo imaging of mutant huntingtin in Huntington’s disease. Sci Transl Med. 2022;14:eabm3682. doi:10.1126/scitranslmed.abm3682.

14. Bertoglio D, Weiss AR, Liguore W, Martin LD, Hobbs T, Templon J, et al. In Vivo Cerebral Imaging of Mutant Huntingtin Aggregates Using (11)C-CHDI-180R PET in a Nonhuman Primate Model of Huntington Disease. J Nucl Med. 2023. doi:10.2967/jnumed.123.265569.

15. Burns HD, Hamill TG, Eng W-s, Hargreaves R. Annual Reports In Medicinal Chemistry - Section VI - Topics in Drug Design and Discovery. United States of America: Academic Press; 2001.

16. Li S, Schmitz A, Lee H, Mach RH. Automation of the Radiosynthesis of Six Different (18)F-labeled radiotracers on the AllinOne. EJNMMI Radiopharm Chem. 2017;1:15. doi:10.1186/s41181-016-0018-0.

17. Liu L, Johnson PD, Prime ME, Khetarpal V, Brown CJ, Anzillotti L, et al. Design and Evaluation of [(18)F]CHDI-650 as a Positron Emission Tomography Ligand to Image Mutant Huntingtin Aggregates. J Med Chem. 2022. doi:10.1021/acs.jmedchem.2c01585.

18. Menalled LB, Kudwa AE, Miller S, Fitzpatrick J, Watson-Johnson J, Keating N, et al. Comprehensive behavioral and molecular characterization of a new knock-in mouse model of Huntington’s disease: zQ175. PLoS One. 2012;7:e49838. doi:10.1371/journal.pone.0049838.

19. Heikkinen T, Lehtimaki K, Vartiainen N, Puolivali J, Hendricks SJ, Glaser JR, et al. Characterization of neurophysiological and behavioral changes, MRI brain volumetry and 1H MRS in zQ175 knock-in mouse model of Huntington’s disease. PLoS One. 2012;7:e50717. doi:10.1371/journal.pone.0050717.

20. Cudalbu C, McLin VA, Lei H, Duarte JM, Rougemont AL, Oldani G, et al. The C57BL/6J mouse exhibits sporadic congenital portosystemic shunts. PLoS One. 2013;8:e69782. doi:10.1371/journal.pone.0069782.

21. Bertoglio D, Verhaeghe J, Korat S, Miranda A, Wyffels L, Stroobants S, et al. In vitro and In vivo Assessment of Suitable Reference Region and Kinetic Modelling for the mGluR1 Radioligand [(11)C]ITDM in Mice. Mol Imaging Biol. 2020;22:854–63. doi:10.1007/s11307-019-01435-1.

22. Bertoglio D, Verhaeghe J, Miranda A, Kertesz I, Cybulska K, Korat S, et al. Validation and noninvasive kinetic modeling of [(11)C]UCB-J PET imaging in mice. J Cereb Blood Flow Metab. 2020;40:1351–62. doi:10.1177/0271678X19864081.

23. Verhaeghe J, Bertoglio D, Kosten L, Thomae D, Verhoye M, Van Der Linden A, et al. Noninvasive Relative Quantification of [(11)C]ABP688 PET Imaging in Mice Versus an Input Function Measured Over an Arteriovenous Shunt. Front Neurol. 2018;9:516. doi:10.3389/fneur.2018.00516.

24. Miranda A, Bertoglio D, Glorie D, Stroobants S, Staelens S, Verhaeghe J. Validation of a spatially variant resolution model for small animal brain PET studies. Biomed Phys Eng Express. 2020;6:045001. doi:10.1088/2057-1976/ab8c13.

25. Bertoglio D, Verhaeghe J, Kosten L, Thomae D, Van der Linden A, Stroobants S, et al. MR-based spatial normalization improves [18F]MNI-659 PET regional quantification and detectability of disease effect in the Q175 mouse model of Huntington’s disease. PLoS One. 2018;13:e0206613. doi:10.1371/journal.pone.0206613.

26. Johnson GA, Badea A, Brandenburg J, Cofer G, Fubara B, Liu S, et al. Waxholm space: an image-based reference for coordinating mouse brain research. Neuroimage. 2010;53:365–72. doi:10.1016/j.neuroimage.2010.06.067.

27. Cicchetti D. Guidlines, Criteria, and Rules of Thum for Evaluating Nomred and Standardized Assessment Instruments in Psychology. Psychological Assessment. 1994;6:284–90. doi:1040-3590/94/53.00.

28. Kaur T, Brooks AF, Lapsys A, Desmond TJ, Stauff J, Arteaga J, et al. Synthesis and Evaluation of a Fluorine-18 Radioligand for Imaging Huntingtin Aggregates by Positron Emission Tomographic Imaging. Frontiers in Neuroscience. 2021;15. doi:10.3389/fnins.2021.766176.

